# Dual checkpoint blockade of glioblastoma with Anti-PD-1 and Anti-LAG-3 promotes expansion of tumor-reactive T cell clones along a unique pathway of differentiation

**DOI:** 10.1101/2025.09.06.674490

**Authors:** Mingshuang Wang, Yuntian Fu, Sadhana Bom, Yi Ning, Divij Matthews, Ming Zhang, Calixto-Hope Lucas, John Choi, Zhen Zeng, Tianbei Zhang, Hongkai Ji, Vasan Yegnasubramanian, Jon Weingart, Chetan Bettegowda, Kellie N. Smith, Michael Lim, Drew Pardoll, Nancy R. Zhang, Christina Jackson

## Abstract

IDH-wildtype grade IV glioblastoma is the most aggressive adult primary brain tumor and remains refractory to anti-PD-1 monotherapy despite evidence of limited tumor-specific T cell induction. To determine the impact of immune checkpoint inhibitors (ICIs) on glioblastoma T cell transcriptional landscape and repertoire, we conducted paired single-cell RNA sequencing (scRNA-seq) and T cell receptor sequencing (TCR-seq) of tumor-infiltrating lymphocytes (TILs) from patients with untreated, newly diagnosed glioblastoma and from recurrent glioblastoma treated with dual checkpoint blockade targeting PD-1 and LAG-3. Using a validated transcriptional signature, we found that predicted tumor-reactive T cells (TRC) in untreated glioblastomas reside almost exclusively in a clonally expanded GZMK^hi^ population with developmental plasticity, affording them the potential to differentiate into both tissue-resident and terminal effector T cells. Dual ICI therapy induced substantial clonal remodeling, characterized by the recruitment of new TRC from the periphery into the tumor microenvironment (TME) and differentiation into transitional effectors and ultimately terminal effectors along a gradient characterized by simultaneous acquisition of cytotoxic and exhaustion genes, regulated by specific transcriptional, metabolic, and epigenetic programs. Longitudinal clonal tracking in peripheral blood confirmed that with ICI treatment, most TRC expand transiently in circulation prior to tumor infiltration, with peripherally derived clones becoming the major contributor to the GZMK^hi^ TRC that further expand in the tumor. Our study provides the first comprehensive map of T cell clonal dynamics and differentiation in glioblastoma following dual ICIs and highlights a potential mechanism of immune activation and peripheral recruitment of TRC in glioblastoma not previously described. Our results suggest that therapeutic strategies to sustain these GZMK^hi^ early effector and transitional effector T cells may further enhance ICI therapeutic efficacy in glioblastoma.

## Main Text

IDH-wildtype grade IV glioblastoma is the most common and aggressive primary malignant brain tumor in adults^1^. Despite maximal surgical resection, radiation, and chemotherapy, glioblastoma is universally fatal, with a median survival of 15 months^2^. Advances in glioblastoma therapy have been marginal in the last two decades; hence, there is a desperate need for novel therapeutic modalities and approaches. Treatment with single immune checkpoint inhibitor (ICI), which has produced durable disease-free response in other malignancies, has largely failed in glioblastoma^2–4^, with the exception of DNA repair deficient tumors^5,6^.

There is broad consensus that limited ICI efficacy in glioblastoma is due to potent immunosuppressive mechanisms that counteract T cell activation including systemic lymphopenia due to sequestration of T cells in the bone marrow, infiltration of regulatory T cells, and a large representation of tumor-associated suppressive myeloid cells^7–9^. In addition to external immunosuppressive barriers, emerging evidence points to critical T cell–intrinsic mechanisms of resistance in glioblastoma. High-dimensional flow cytometric analysis in murine models of glioblastoma reveal dysfunctional profiles in glioblastoma-infiltrating T cells, characterized by the co-expression of multiple immune checkpoint molecules (PD-1, TIM-3, LAG-3) and upregulation of transcriptional factors such as TOX^10,11^. However, single-cell RNA sequencing (scRNA-seq) of human glioblastoma tumor-infiltrating CD8^+^ T cells (TILs) has shown conflicting results, lacking a consistent transcriptomic and proteomic profile attributed to exhausted T cells in both IDH-wildtype and IDH-mutant gliomas^12,13^. These contrasting observations underscore the complexity of T cell dysfunction in glioblastoma; therefore, a comprehensive and integrative approach is necessary to resolve these contradictions and fully understand the underlying mechanisms of T cell dysfunction in glioblastoma.

Despite lack of efficacy with ICI in glioblastoma, there is evidence that ICI does impact the glioblastoma TME with neoadjuvant administration of ICI promoting survival benefits in a subset of patients with promotion of at least some tumor-specific T cells^14–17^. Furthermore, given evidence of non-redundant mechanisms of T cell dysfunction conferred by different checkpoints^18–20^, there is increasing interest in combining multiple ICIs for added efficacy^18,21,22^. In murine glioblastoma models, combination checkpoint inhibition with anti-PD-1 and anti-LAG-3 led to improved survival^23^. This finding was recapitulated in our clinical trial (ABTC 1501)- the first study to evaluate the safety and efficacy of combined anti-PD1 and anti-LAG-3 therapy in patients with recurrent glioblastoma-which demonstrated enhanced CD8^+^ T cell infiltration and durable responses in a significant subset of patients. However, the impact of dual ICIs on glioblastoma T cell landscape and repertoire remains unknown.

In the present study, leveraging patient-derived samples from our clinical trial, we performed scRNA-seq coupled with paired T cell receptor (TCR) sequencing (TCR-seq) on surgically resected tumor samples from various untreated adult diffuse glioma subtypes, including 21 treatment-naïve glioblastoma patients and 5 glioblastoma patients treated with combination anti-PD-1 and anti-LAG-3 therapy. Analyzing the largest single-cell multiomic brain cancer TIL dataset, we aimed to comprehensively dissect the T cell landscape in glioblastoma, with particular attention to the impact of combination ICIs on glioblastoma TME and T cell landscape to elucidate determinants of immunotherapy response and resistance.

Integrating novel methodologies to identify tumor-reactive T cells (TRC) by transcriptional signature and analyzing clonal T cell dynamics using innovative computational approaches, we find that the most clonally expanded population of T cells in glioblastoma consisted of GZMK^+^ T cells with developmental plasticity that are capable of diverging into multiple distinct states. Our results revealed a paucity of TRC in untreated glioblastoma patients. However, treatment with dual ICIs recruits new TRC from the periphery and drives GZMK^+^ effector memory T cells to proliferate and differentiate along an effector trajectory that initially upregulates a cytotoxic program but ultimately develops an exhaustion program, characterized by upregulation of TCR signaling, proliferation, TOX2 expression, and checkpoint expression. Our work provides key mechanistic insights into how dual ICIs reshape the T cell landscape and repertoire in glioblastoma and identifies potential mechanisms of immunotherapy resistance in these patients, highlighting opportunities for therapeutic optimization.

## Results

### Tumor-reactive T cells in glioblastoma exhibit GZMK-high CD8 effector phenotypes

To define the transcriptional states and clonal diversity of TILs in glioblastoma, we performed droplet-based 5′ scRNA-seq with paired TCR-seq on freshly isolated CD45⁺CD3⁺ T cells obtained by fluorescence-activated cell sorting (FACS). Samples included six surgically resected IDH-mutant, 1p/19q-codeleted grade 2 oligodendrogliomas; one IDH-mutant grade 2 astrocytoma; three IDH-mutant grade 3 astrocytomas; two IDH-mutant grade 4 astrocytomas; 21 treatment-naïve IDH-wildtype glioblastomas; and five IDH-wildtype glioblastomas treated with dual anti-PD-1/anti-LAG-3 blockade. Across all tumors, we profiled 129,465 CD8⁺ T cells and 163,561 CD4⁺ T cells (Fig. 1a, Extended Data Table 1).

**Fig. 1.**
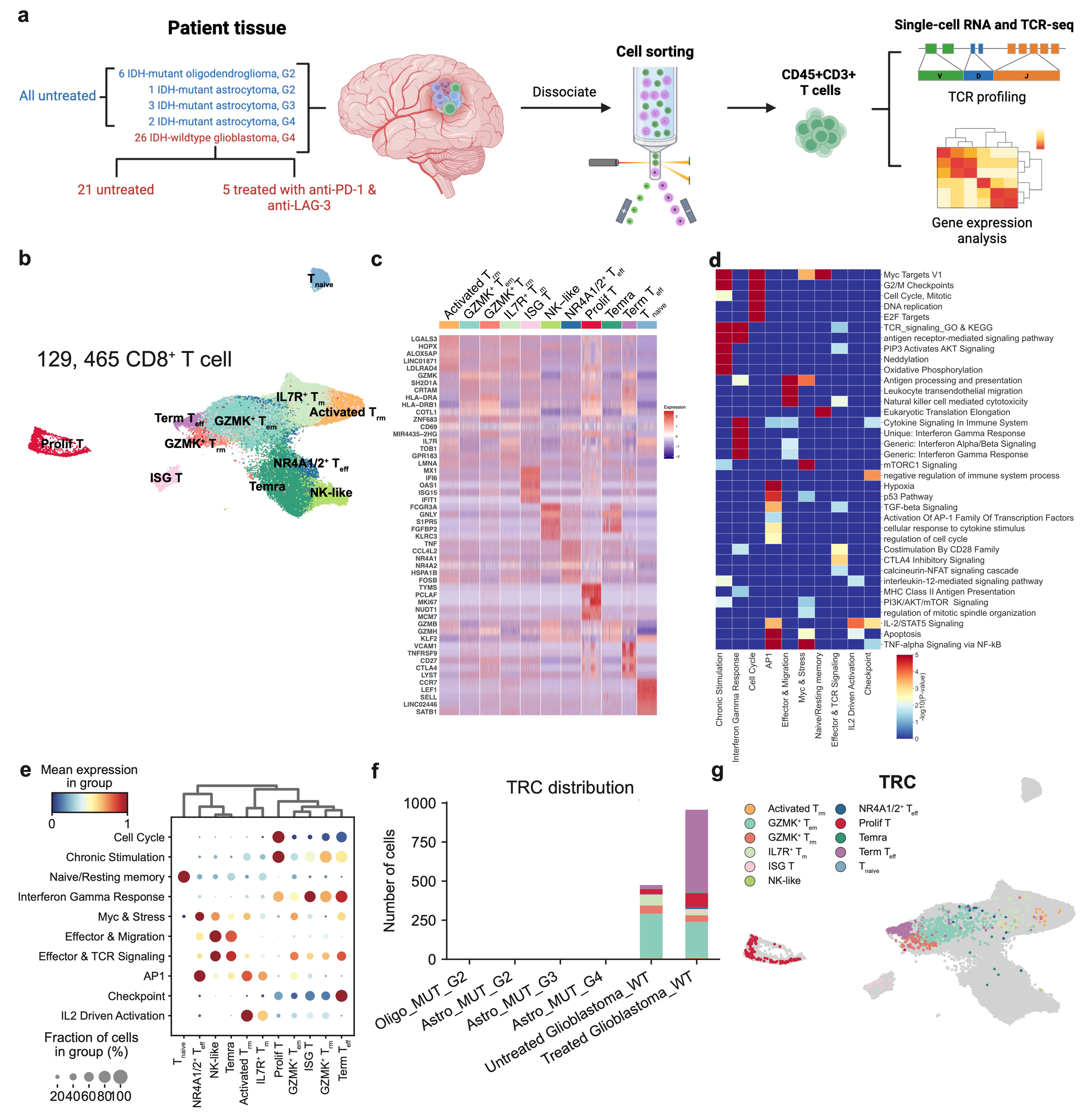
Tumor-reactive T cells in glioblastoma exhibit GZMK^hi^ CD8^+^ effector phenotypes. (a). Schematic diagram showing experimental design. Paired scRNA-seq and TCR-seq are performed on CD45^+^CD3^+^ cells isolated from resected untreated IDH-mutant grade 2 oligodendroglioma (n=6), IDH-mutant grade 2 astrocytoma (n=1), IDH-mutant grade 3 astrocytoma (n=3), IDH-mutant grade 4 astrocytoma (n=2), IDH-wildtype glioblastoma (n=21), and dual immune checkpoint inhibitor-treated IDH-wildtype glioblastoma (n=5). (b). UMAP projection of the expression profile of CD8÷ T cells (not including MAITT cells) in gliomas demonstrating 11 clusters of CD8^+^ T cells, each delineated by a color code. (c). Z-score-normalized heatmap of the top five differentially expressed genes with the highest log fold-change from each CD8^+^ T cell cluster, (d). Heatmap demonstrating enrichment of specific transcriptomic pathways across each correlated functional module identified using Hotspot, (e). Dotplot displaying differences in expression of functional modules across clusters of CD8^+^ T cells demonstrates that GZMK^+^ T_em_ cells exhibit a broad but moderate expression across multiple gene modules, (f). Distribution of tumor-reactive T cells (TRC) predicted based on two cells within a clone that are MANAscore-high demonstrates that TRC are exclusively present in IDH-wildtype glioblastoma and are further increased in treated patient samples, (g). UMAP demonstrates that the majority of TRC reside in Term T_eff_, GZMK^+^ T_em_, GZMK÷T_rm_, and Prolif T cells.

Unbiased clustering of transcriptional profiles revealed 11 CD8^+^ T cell clusters, 11 CD4^+^ T cell clusters, as well as discrete subsets of MAIT and γδT cells (Extended Data Fig. 1a, Extended Data Table 2). Given their role as principal cytotoxic effectors and key mediators of ICI responses, we focused subsequent analyses on the CD8⁺ compartment. Within the CD8⁺ compartment, we identified clusters corresponding to canonical T cell states, including naive T cells (T_naive_), central memory T cells (IL7R^+^ T_m_), terminally differentiated effector memory T cells re-expressing CD45RA (Temra), effector T cells (T_eff_), resident memory T cells (T_rm_), effector memory T cells (T_em_), and NK-like T cells (NK-like) (Fig. 1b-c). Consistent with recent observations by Wang et al.^13^ that a large proportion of overall glioblastoma TILs exhibited high *GZMK* expression, in our dataset, this was particularly pronounced in the T_em_, T_rm_, Term T_eff_, interferon-stimulated gene-high (ISG T), and proliferating (Prolif T) T cell clusters (Extended Data Fig. 1b).

Using Hotspot^24^ and Gene Set Enrichment Analysis (GSEA), we delineated ten transcriptional modules representing discrete functional programs (Fig. 1d-e, Extended Data Fig. 1c)^25^. These modules captured intersecting but non-redundant pathways underlying CD8⁺ T cell heterogeneity in the glioblastoma tumor microenvironment (TME). Among them, we identified transcriptional programs corresponding to canonical functions such as Chronic Stimulation, Interferon Gamma Response, Effector & TCR Signaling, and Checkpoint regulation. Distinct clusters showed preferential enrichment for specific programs. For example, Term T_eff_ cells were enriched in Interferon Gamma Response, Chronic Stimulation, and Checkpoint modules, consistent with sustained TCR engagement and features of functional exhaustion^26,27^. Prolif T cells were enriched in Cell Cycle, Chronic Stimulation, and Interferon Gamma Response modules, reflecting a cytokine-responsive, proliferative phenotype adapted to chronic antigen exposure. ISG T cells, as expected, were enriched for interferon-related programs. Notably, deconvolution of these signatures revealed simultaneous activation of type I (IFN-α/β) and type II (IFN-γ) interferon response pathways, with a modest bias toward type I signaling, suggesting localization to a distinct intratumoral niche shaped by persistent tumor-derived or microenvironmental innate pathogen-associated molecular pattern (PAMP) or damage-associated molecular pattern (DAMP)-derived signals (Extended Data Fig. 1d-e)^28^. Finally, while both Effector & Migration and Effector & TCR Signaling modules encoded cytotoxic machinery, their expression patterns varied across subsets. NK-like and Temra cells displayed broad, nonselective expression of both programs. By contrast, GZMK^+^ T_em_ and Term T_eff_ subsets selectively engaged the tightly regulated Effector & TCR Signaling program, reflecting a more controlled mode of cytotoxic activation^29^.

Among the intratumoral CD8^+^ T cell states, the GZMK⁺ T_em_ population stands out as a uniquely transcriptionally plastic subset. Unlike other clusters that are dominated by specific transcriptional programs, GZMK⁺ T_em_ cells exhibit a broad but moderate expression across multiple gene modules, without exclusive enrichment for any single program. This includes partial engagement of Effector & TCR Signaling, Effector & Migration, AP1, Myc & Stress, and Checkpoint modules, suggesting that these cells straddle multiple functional axes—effector potential, metabolic readiness, inflammatory response, and regulatory control. This diffuse modular profile implies that GZMK⁺ T_em_ cells represent a plastic population within the CD8⁺ T cell differentiation continuum—sharing memory-like, effector, and exhaustion features. Their modest expression of checkpoint molecules indicates recent or intermediate antigen experience without full terminal exhaustion. Simultaneously, their enrichment in effector modules, including cytotoxic and chemotactic genes, points to latent functional capacity. The co-expression of Myc & Stress and AP1 modules further supports the notion that these cells are in a metabolically primed and environmentally responsive state, potentially regulated by dynamic external cues within the TME. Crucially, the GZMK⁺ T_em_ population is one of the few clusters that simultaneously expresses genes involved in early activation (IL2 signaling, TCR responsiveness) and immune regulation (checkpoints, stress response). Together, these results reveal that CD8⁺ T cells in glioblastoma span a spectrum of activation, functional specialization, and exhaustion states. The GZMK⁺ T_em_ population emerges as a central and transcriptionally pliable subset, enriched for overlapping modules but not dominated by any single one, suggesting lineage plasticity and therapeutic potential. In contrast, terminal and chronically stimulated subsets are defined by concentrated expression of cell cycle, checkpoint, and metabolic modules, pointing toward functional rigidity and exhaustion.

Consistent with their relatively low mutational burden, glioblastomas generally harbor sparse TIL populations within the TME. Beyond limited reported cases of mutation-associated neoantigen (MANA)-specific TCR identified in neoantigen-vaccinated patients^14,30^, little is known regarding the frequency of truly tumor-specific CD8^+^ TILs or their associated transcriptional states. To begin addressing this question, we first examined potential categories of antigen specificity within our TIL dataset. We leveraged previously published gene signatures derived from CD8^+^ TILs across multiple cancer types, in which TCR had been experimentally validated to recognize either viral antigens (i.e. influenza, EBV) or tumor-specific antigens (Extended Data Fig. 1f, Extended Data Table 3)^31,32^. We further employed our recently developed three gene MANAscore algorithm to more accurately identify predicted TRC among our CD8^+^ TILs^33^. This algorithm, developed using ensembled single-patient voting models, calculates a MANAscore from both imputed and non-imputed mRNA expression of *CXCL13, CD39,* and *IL7R*. MANAscore-high cells were then defined independently for each patient, with cutoffs determined from the distributions of the imputed and non-imputed MANAscores. TRC are subsequently inferred as cells within those clones that share the same TCR as the MANAscore-high cells. This approach was shown to accurately predict TRC in TILs specific for MANAs, tumor associated antigens, and tumor viral antigens across more than five human tumor types^33^. We observed that MANAscore-high TILs within glioblastomas localized predominantly within the Term T_eff_ population (Extended Data Fig. 1g-h). Predicted TRC were identified as clonotypes containing at least one MANAscore-high cell and represented by at least two cells within a given patient. Using this approach, we successfully identified TRC despite their representing a small proportion of total CD8^+^ TILs in glioblastoma. Notably, there were essentially no MANAscore-high cells, and thus no TRC, in IDH-mutant gliomas (Fig. 1f, Extended Data Fig. 1g). The vast majority of predicted TRC in glioblastoma were confined to four transcriptional states: Term T_eff,_ GZMK^+^ T_em_, GZMK^+^ T_rm_, and Prolif T cells, but with a significant shift from GZMK^+^ T_em_ toward Term T_eff_ upon ICI treatment (Fig. 1f-g).

To correlate whether CD8^+^ T cell clusters with the highest enrichment of TRC also exhibited greater clonal expansion from antigen stimulation, we employed a novel frequency-based expansion index to measure the clonal size within each cluster normalized by cluster size across all CD8^+^ T cell subsets (see Methods). We found correspondingly that the GZMK^+^ T_em_ population is the most clonally expanded population in untreated glioblastoma (Extended Data Fig. 2a). In addition, GZMK^+^ T_em_ cells in IDH-wildtype glioblastomas exhibited a distinct transcriptomic profile compared to those in other subtypes, including IDH-mutant grade 4 astrocytomas (Fig. 2a). In glioblastomas, these GZMK^+^ T_em_ cells showed increased signatures of T cell activation and proliferation relative to IDH-mutant gliomas, suggesting a potential critical role for this population in shaping the glioblastoma TME (Fig. 2b). Notably, upon treatment with dual ICIs, both Term T_eff_ and GZMK^+^ T_em_ populations demonstrated the most significant increase in both abundance and clonal expansion, with GZMK^+^ T_em_ cells exhibiting the greatest degree of expansion (Fig. 2c-d). These findings suggest that GZMK^+^ T_em_ and Term T_eff_ cells are the primary responding cells to immunotherapy in glioblastoma, suggesting that they may play an important role in determining responsiveness to therapy. Strikingly, we found that in untreated glioblastoma, the vast majority of the TRC resided in the GZMK^+^ T_em_ population and were absent from the Term T_eff_ population. Following dual checkpoint blockade, however, TRC shifted predominantly to the Term T_eff_ population (Fig. 2e). These findings suggest a model in which checkpoint inhibition induces differentiation from a progenitor GZMK^+^ T_em_ population ultimately toward a Term T_eff_ population. We next sought to validate this trajectory at the clonal level.

**Fig. 2.**
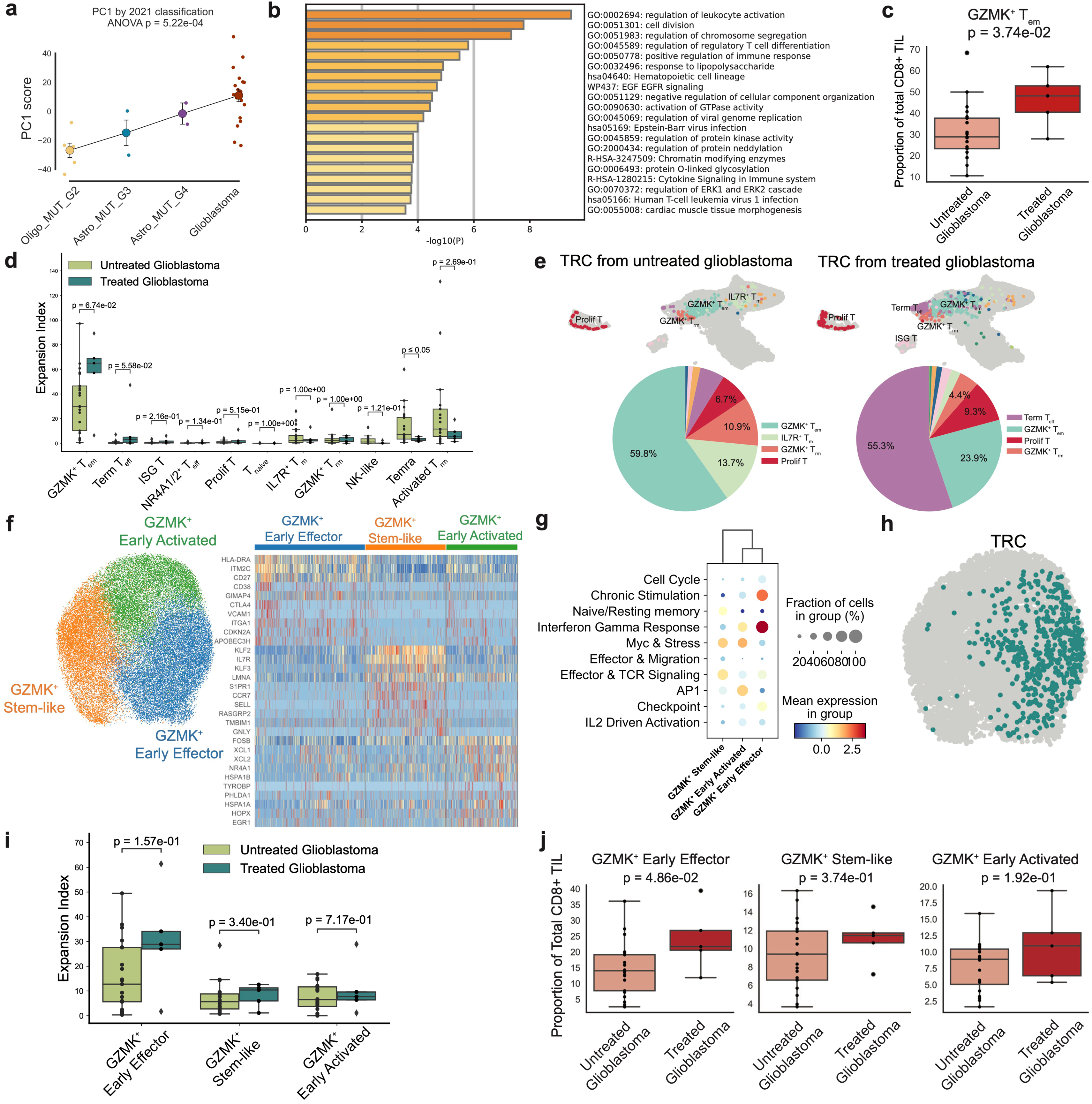
Fine structure analysis of GZMK^+^ T_em_ reveals multiple transcriptional programs associated with tumor grade, tumor specificity, and ICI treatment. (a). Pseudobulk principal component analysis of the GZMK^+^ T_em_ cells demonstrates marked differences in transcriptomic profiles of these cells in glioblastomas compared to those in other subtypes of gliomas, (b). Gene set enrichment analysis demonstrates that GZMK^+^ T_em_ cells in glioblastoma show increased signatures of T cell activation and proliferation relative to GZMK^+^ T_em_ from IDH-mutant gliomas, (c). Boxplot demonstrating increased infiltration of GZMK^+^ T_em_ in treated glioblastoma patients versus treatment-naive patients, (d). Expansion index analysis shows that GZMK^+^ T_em_ and Term T_eff_ cells have the highest change in clonal expansion after treatment with dual ICIs. (e). Identification of TRC in untreated and treated patients shows enrichment of TRC in the GZMK^+^ T_em_ population in untreated patients (left) versus enrichment of TRC in Term T_eff_ in treated patients (right), (f). Sub-clustering of GZMK^+^ T_em_ reveals three subsets with distinct cellular states. Heatmap demonstrating the top 10 differentially expressed genes in each subset leading to the identification of GZMK^+^ early effector, GZMK^+^ stem-like, and GZMK^+^early activated subsets, (g). Dotplot demonstrating relative expression of functional Hotspot modules across GZMK^+^ subsets shows enrichment of chronic stimulation and IFNy modules in GZMK^+^ early effector cells, (h). Evaluation of previously identified TRC demonstrates the exclusive presence of TRC in GZMK^+^ early effector subset, (i). Expansion index comparison shows that GZMK^+^ early effector cells demonstrate the most significant increase in clonal expansion after treatment, (j). Boxplot showing selective increase in infiltration of GZMK^+^ early effector cells after treatment with dual ICIs.

### Dual checkpoint blockade preferentially expands an early effector subset of GZMK^+^ T_em_ cells

To further dissect intratumoral differentiation pathways, we next focused specifically on the GZMK^+^ T_em_ CD8^+^ population. As demonstrated by module analysis, the GZMK^+^ T_em_ cluster displays an intermediate expression of multiple modules, suggesting that it is a potentially dynamic and heterogeneous cell cluster (Fig. 1e). Therefore, we further sub-clustered the GZMK^+^ T_em_ population and identified three distinct cellular states (Fig. 2f). The GZMK^+^ early effector subset co-expresses genes encoding activation antigens (*CD27, HLA-DRA, CD38*) alongside T cell checkpoints (*CTLA4*). The GZMK^+^ stem-like subset expresses genes encoding classical memory and naive markers (*CCR7, IL7R, SELL*). Finally, the GZMK^+^ early activated subset is characterized by early activation gene signatures, including *FOSB, EGR1, XCL1,* and *XCL2*. While all clusters demonstrated expression of *GZMK*, their functional programs diverged. Genes associated with cytotoxicity (*GZMB, GNLY, and GZMA)* are preferentially expressed in the GZMK^+^ early effector subset, genes indicative of stemness (*TCF7*) are expressed in the GZMK^+^ stem-like subset, and genes for early T cell activation are enriched in the GZMK^+^ early activated subset (Extended Data Fig. 2b). Transcription factor (TF) analysis further reinforced these distinctions. GZMK^+^ early effector cells showed enhanced activity of regulons that play crucial roles in T cell effector function (EOMES, STAT1). Correspondingly, GZMK^+^ stem-like cells exhibited upregulation of Kruppel-like factors (KLF3, KLF2), linked to T cell quiescence in naïve and resting memory states. Lastly, GZMK^+^ early activated cells demonstrated strong activation of TFs associated with the AP-1 pathway and early-stage activation (FOSB, EGR1) (Extended Data Fig. 2c).

Our initial module analysis (Fig. 1e) revealed that these GZMK^+^ T_em_ cells span a spectrum of transcriptomic states. Further stratification revealed that these GZMK^+^ subsets map onto distinct functional modules, with the GZMK^+^ early effector cells demonstrating strong upregulation of interferon-γ response, chronic stimulation, and checkpoint programs (Fig. 2g). Notably, within the GZMK^+^ T_em_ subclusters, TRC localized exclusively within the GZMK^+^ early effector subset (Fig. 2h), which also exhibited the highest clonal expansion and increased markedly in frequency following dual immune checkpoint blockade (Fig. 2i-j). These findings suggest that the GZMK^+^ early effector subset could serve as the key precursor population that gives rise to Term T_eff_ cells in response to treatment.

To dissect differentiation trajectories within GZMK+ T_em_ cells at the clonal level, as well as the influence of immunotherapy upon these processes, we applied Clonotrace, a novel lineage tracing–based algorithm designed to resolve cell dynamics. Clonotrace groups T cell clones with similar cell state distributions into distinct clonotype profiles, where a clonotype profile is a set of clonotypes that traverse the same transcriptomic states (Fig. 3a left). By integrating the clonotype profile embedding with transcriptional information, Clonotrace distinguishes not only transcriptional similarity but also lineage relationships, such that cells with similar gene expression but divergent clonal profiles are separated farther in two-dimensional space. This results in a modified UMAP that enhances the resolution of differentiation branches and enables identification of fate-driving genes (Fig. 3a right). Overall, Clonotrace identified 9 distinct clonotype profiles across CD8^+^ T cells (Fig. 3a left). Although the Prolif T cluster was included in constructing the clonal embedding, it was excluded from the weighted transcriptional UMAP and pseudotime analyses because its strong cell cycle signature obscured the underlying differentiation trajectory. Of the 9 clonotype profiles, profiles 4, 5, 6, and 9 were restricted to a single cell state, whereas profiles 1, 2, 3, 7, and 8 spanned multiple states, enabling inference of clonal differentiation trajectories (Extended Data Fig. 3a). Among these, profiles 1, 2, and 3 traverse the GZMK^+^ early effector (GZMK^+^ T_em_) cluster, suggesting dynamic transitions within this population. Of these three profiles, profile 3 was observed in all gliomas while profiles 1 and 2 are highly enriched in glioblastomas (Fig. 3b, Extended Data Fig. 3b). To define the directionality of differentiation within these profiles, we examined the T cell repertoire from a patient with paired scRNA/TCR-seq data obtained at two consecutive surgical timepoints during treatment. At timepoint 2, we observed marked expansion of GZMK^+^ early effector, Term T_eff_, ISG T, and GZMK^+^ T_rm_ populations relative to other subsets (Extended Data Fig. 3c). To further dissect clonal relationships, we constructed a TCR clonal sharing matrix across these four T cell states and quantified the relative contribution of expanded cellular states at timepoint 2 by those present at timepoint 1. Clonal sharing analyses revealed that GZMK^+^ early effector cells were the major contributors to ISG T cells in Profile 1 and to Term T_eff_ cells in Profile 2 suggesting GZMK^+^ early effectors as the earlier cellular state (Extended Data Fig. 3d). To further support this inferred trajectory, we calculated exhaustion scores amongst these subsets using signatures from Giles et al^34^. Analogous to T cell progression to exhaustion in the setting of chronic viral infection, exhaustion scores increased progressively from the GZMK^+^ early effector population to Term T_eff_ cells (Extended Data Fig. 3e). Together, these findings suggest that GZMK^+^ early effector cells can follow two potential differentiation trajectories: (1) toward T_rm_/ISG T cell fates (trajectory 1) and (2) toward terminal effector T cell (Term T_eff_) fate (trajectory 2) (Fig. 3c). Notably, trajectory 1 (Profile 1) clones predominated in untreated glioblastomas, whereas trajectory 2 (Profile 2) clones were selectively expanded in ICI-treated glioblastoma patients, indicating that ICI drives GZMK^+^ early effectors preferentially toward terminal effector differentiation.

**Fig. 3.**
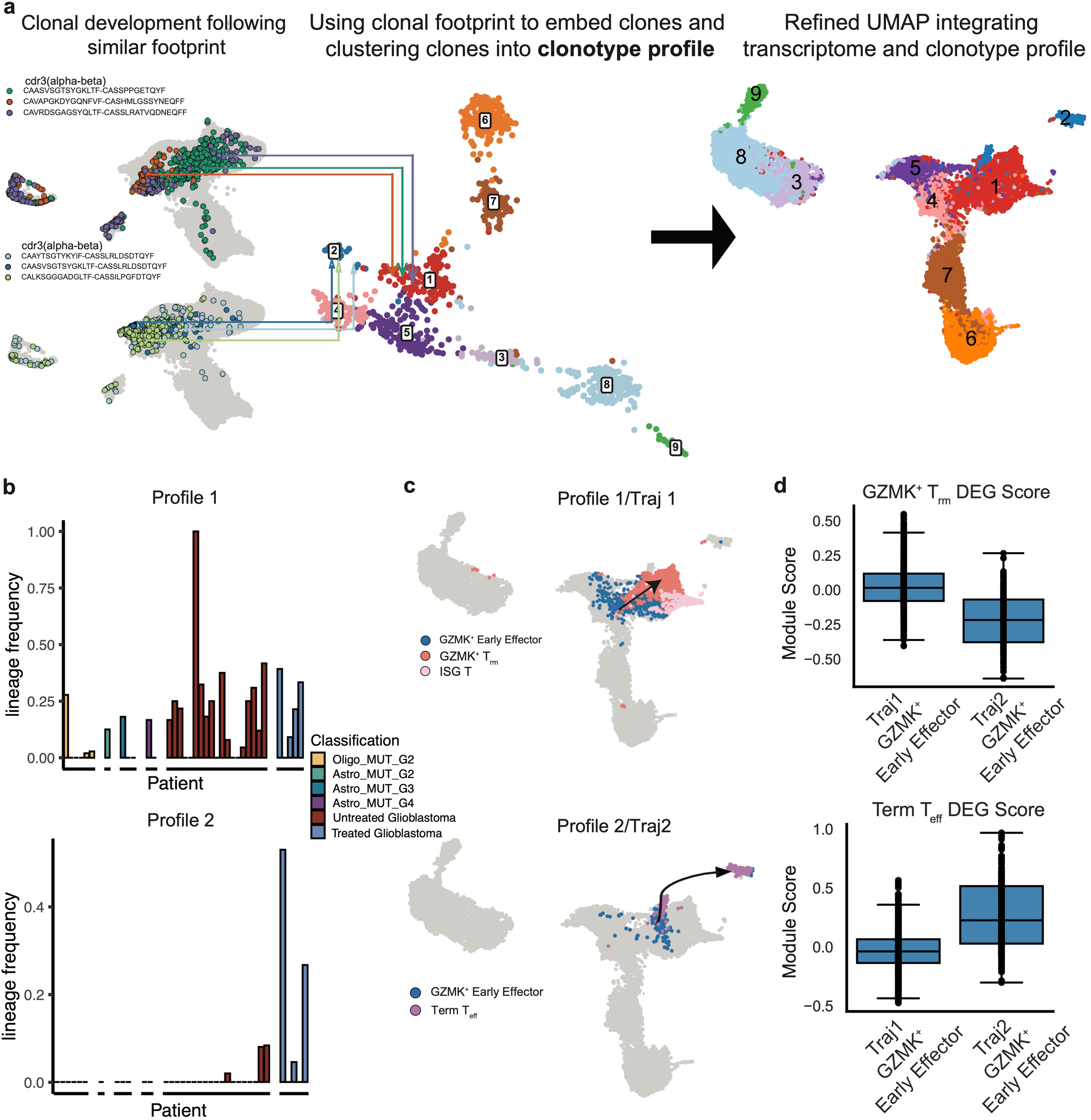
Clonotrace reveals bifurcated clonal dynamics of GZMK^+^ early effector cells stratified by ICI treatment status. (a). Clonotrace identifies nine clonotype profiles based on similarity in transcriptomic footprint (left). Integration of clonotype profile and transcriptomic information results in modified UMAP that enhances the resolution of cellular dynamics and differentiation (right). Prolif T cells are included in constructing the clonal embedding but excluded from the clonal embedding weighted transcriptional UMAP and pseudotime analysis because their strong cell cycle signature obscures the underlying differentiation trajectory, (b). Bar plot demonstrating the enrichment of clonotype profiles 1 and 2 in glioblastoma patients, with each bar representing one patient. Clonotype profile 2 is specifically enriched in ICI-treated glioblastoma patients, (c). Comparison of shared clones at two time points of treatment within a patient demonstrates bifurcated differentiation trajectories centered at GZMK^+^ early effector cells toward either resident-memory cells (GZMK^+^ T_rm_) or terminal effector cells (Term T_eff_). (d). Comparison of differentially expressed genes between precursor populations (GZMK^+^ early effectors in trajectory 1 or trajectory 2) and terminal cells (GZMK^+^ T_rm_ or Term T_eff_) demonstrates similar upregulation of transcriptional signatures in the precursor states to their terminal counterparts along these two trajectories.

To investigate the molecular determinants of this bifurcation, we compared the differences in transcriptomic programs between GZMK^+^ early effector subsets that are restricted to the two trajectories. GZMK^+^ early effectors that are committed to trajectory 2, which are expanded by ICI treatment, exhibited transcriptional signatures resembling their terminal state counterpart (Term T_eff_) even at an early stage, including upregulation of genes associated with T cell activation (*TNFRSF9, LAG3, STAT3*). In contrast, the GZMK^+^ early effector subset in trajectory 1 demonstrated stronger enrichment of transcriptional signatures similar to GZMK+ T_rm_ cells (*GZMH, ZNF683*), the terminal cellular state in trajectory 1 (Fig. 3d, Extended Data Fig. 3f). These results indicate that changes in trajectory-specific gene programs occur early in the GZMK^+^ early effectors along their respective differentiation paths. Collectively, these data demonstrate that ICI reshapes the developmental trajectory of GZMK^+^ T_em_ cells by directing their GZMK^+^ early effector precursors toward a terminal effector and dysfunctional fate. This suggests that the therapeutic efficacy, and eventual exhaustion, of CD8^+^ T cells in glioblastoma may be determined early during ICI response and shaped by treatment-induced reprogramming within a plastic effector precursor pool.

### Dual checkpoint blockade reprograms tumor-reactive T cells toward an effector and dysfunctional trajectory along an exhaustion gradient

As shown above, TRC in untreated glioblastomas are largely confined to the GZMK^+^ T_em_ compartment, specifically within the GZMK^+^ early effector subset. To understand how dual ICI influences their differentiation, specifically along trajectories 1 and 2, we analyzed gene signatures across these Clonotrace-defined trajectories for the predicted TRC. Notably, we found that tumor-reactive gene programs and TRC were highly enriched in trajectory 2 (>70%) and largely absent from trajectory 1 (<5%), indicating their preferential differentiation down this specific developmental pathway (Fig. 4a, Extended Data Fig. 4a). Interestingly, in ICI-native tumors, TRC remained largely arrested in trajectory 1, failing to progress beyond the GZMK^+^ early effector state. In contrast, dual ICI treatment shifted TRC exclusively into trajectory 2, where they underwent progressive differentiation toward terminal effector (Term T_eff_) states marked by high checkpoint expression (Fig. 4b, Extended Data Fig. 4b). These data suggest that dual ICI not only expands TRC but actively redirects their differentiation toward a terminal effector fate (Term T_eff_).

**Fig. 4.**
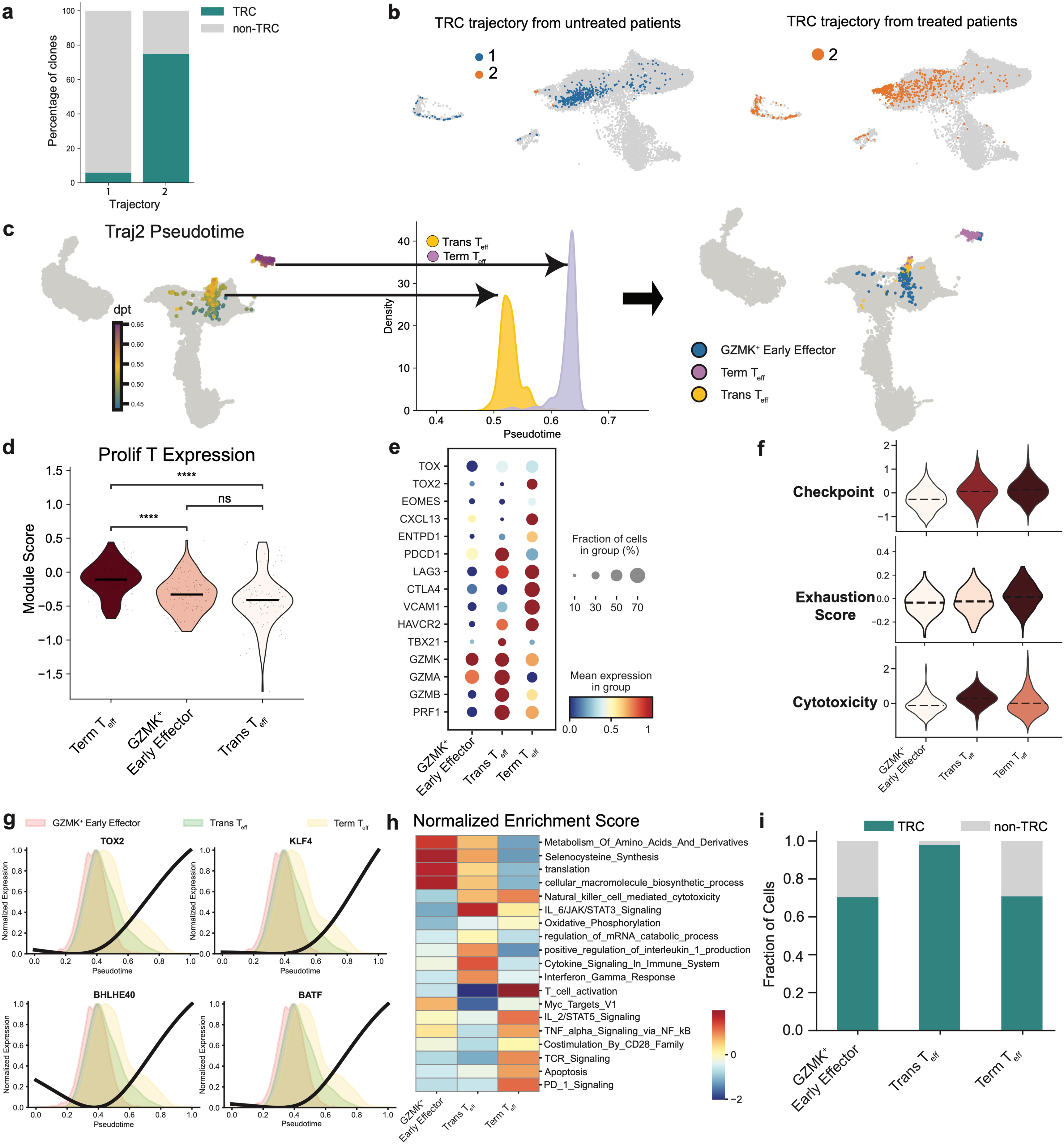
Dual checkpoint blockade reprograms tumor-reactive T cells toward an effector and dysfunctional trajectory along a cytotoxic and exhaustion gradient. (a). Stacked bar plot showing strong enrichment of TRC in trajectory 2 from GZMK^+^ early effector to Term T_eff_. (b). Treatment with dual ICIs drives TRC from trajectory 1 to trajectory 2. (c). Pseudotime analysis of trajectory 2 reveals an intermediate population of transitional effectors (Trans T_eff_) between GZMK^+^ early effector and Term T_eff_ cells, (d). Violin plot demonstrating that Term T_eff_ cells exhibit the highest enrichment of Proliferating T cell transcriptomic signature, (e). Dotplot showing increased expression of cytotoxic cytokines in Trans T_eff_ and increased expression of checkpoints in Term T_eff_ cells, (f). Violin plot showing increasing exhaustion and checkpoint scores as cells progress along trajectory 2 (top), yet Trans T_eff_ demonstrate the highest cytotoxicity score while maintaining low exhaustion score (bottom), (g). Expression of regulons governing effector differentiation and dysfunction along trajectory 2 pseudotime. (h). Heatmap of GSEA-enriched pathways among GZMK^+^ early effector, Trans T_θff_, and Term T_eff_ cells demonstrating upregulation of metabolic and T cell activation pathways in Trans T_eff_. (i). Stacked bar chart showing the highest enrichment of TRC in Trans T_eff_ along trajectory 2.

To further resolve the cellular transitions within trajectory 2, we investigated whether the transition from GZMK+ early effectors to Term T_eff_ cells includes intermediate populations of therapeutic relevance. Although Term T_eff_ cells initially appeared as a single transcriptional cluster in the original UMAP space, Clonotrace trajectory mapping revealed that this subset resolved into two transcriptionally distinct effector subsets (Fig. 4c). Clonal level pseudotime analysis positioned the former as a transitional effector (Trans T_eff_) state, marked by high cytotoxicity and immune responsiveness, while the latter exhibited features of terminal exhausted state. Cell level pseudotime analysis confirmed that Trans T_eff_ cells temporally precede terminal differentiation along trajectory 2, identifying a previously undescribed intermediate state potentially amenable to reinvigoration (Extended Data Fig. 4d).

We next characterized the four key CD8⁺ T cell populations along trajectory 2 - GZMK⁺ early effector, Prolif T, Trans T_eff_, and Term T_eff_, revealing a continuum of transcriptional and functional changes. To determine the proliferative capacity of the three Clonotrace-defined cell states along ICI induced trajectory 2 (GZMK⁺ early effector, Trans T_eff_, and Term T_eff_), we analyzed the enrichment of gene signatures of these three cell states within Prolif T cells. We found that while Prolif T cells contain gene programs from all three states, they most strongly resembled Term T_eff_ cells, suggesting that while proliferation occurs across trajectory 2, but the highest proliferative potential coincides with the transition to the terminal effector/exhaustion state (Fig. 4d). Functionally, GZMK^+^ early effectors expressed minimal exhaustion or cytotoxic markers; Trans T_eff_ cells exhibited peak expression of cytotoxic effector genes (*GZMA, GZMB, PRF1*) and IFN-associated TFs (*STAT1, IRF1, TBX21*); and Term T_eff_ cells upregulated inhibitory receptors (*PDCD1, LAG3, HAVCR2*) and exhaustion-associated TFs (EOMES, TOX2) (Fig. 4e). While checkpoint expression increased progressively across the trajectory, reaching its highest level in Term T_eff_ cells, cytotoxicity peaked earlier in the Trans T_eff_ cells, which simultaneously maintained low exhaustion scores. This pattern suggest that the Trans T_eff_ subset represents a transient window of maximal effector potential (Fig. 4f). Together, these data support a model in which dual ICI therapy reprograms TRC along a unidirectional differentiation path through a transitional state and ultimately toward a dysfunctional end-state. GZMK⁺ early effector cells represent an early plastic state, while Transitional T_eff_ cells emerge as a peak effector population with partial responsiveness. This is followed by cellular proliferation and acquisition of inhibitory receptors and loss of cytotoxic function in Term T_eff_ cells. Therapeutic strategies aimed at stabilizing the Trans T_eff_ state or preventing its transition to exhaustion may enhance anti-tumor responses.

To investigate the molecular underpinnings of this transition, we applied Palantir to infer pseudotime trajectories from the Harmony-integrated manifold and performed trajectory differential gene expression analysis^35^. Prolif T cells, which are dominated by cell cycle-associated genes and presenting a heterogeneous mix of other subsets were excluded from the trajectory analysis, which focused on the progression of T cell effector programs expressed by the GZMK^+^ early effector, Trans T_eff_, and Term T_eff_ populations. This analysis revealed dynamic, stage-specific regulation of TFs governing effector differentiation and dysfunction. Key TFs displayed coordinated yet temporally distinct expression patterns. BATF, a critical effector-associated TF, remained stable early but increased during late pseudotime, potentially sustaining partial effector function under chronic stimulation^36^. BHLHE40, a basic helix-loop-helix transcriptional checkpoint regulator, showed steadily increasing expression across pseudotime. BHLHE40 has been reported as a regulator of virus-specific T cell progression from progenitor to exhausted states in the setting of chronic viral infection^37^ and as an essential mediator of ICI-induced “re-invigoration” of CD8^+^ TILs in murine cancer models^38^. Terminal states along pseudotime demonstrated increased TOX2 and KLF4 expression, which are key regulators of chronic stimulation and terminal effector differentiation, marking the onset of exhaustion-associated programs ^39–42^ (Fig. 4g). Together, these transcriptional transitions delineate an orchestrated shift from early plasticity and effector function to terminal dysfunction, highlighting key intervention points along the TRC differentiation axis.

To further define the functional properties of the Trans T_eff_ population, we performed GSEA across the four populations in trajectory 2. We found that Trans T_eff_ cells and their precursor, GZMK^+^ early effectors, were enriched for gene sets related to cytokine signaling, amino acid metabolism, and IFN-γ pathways (Fig. 4h), consistent with a metabolically and transcriptionally active effector state. Notably, TRC were most abundant in the Trans T_eff_ compartment, underscoring its importance as a key anti-tumor effector population (Fig. 4i). Collectively, our findings demonstrate that dual PD-1/LAG-3 blockade induces a distinct transcriptional trajectory of TRC differentiation defined by early activation, transient cytotoxicity, and eventual exhaustion. The emergence of a metabolically and transcriptionally active Trans T_eff_ population suggests a narrow but actionable therapeutic window in which reinvigoration strategies aimed at sustaining effector function may be most effective.

### Dual ICIs leads to recruitment of new tumor-reactive cells from the periphery into the tumor microenvironment

In cancers such as basal cell carcinoma, melanoma, and non-small cell lung cancer, checkpoint blockade has been shown to reshape the TIL repertoire by both expanding pre-existing T cell clones and recruiting new, peripherally derived clones into the TME^15,43,44^. However, these studies did not determine whether the recruited clones were tumor-specific. As such, the dynamics of TRC in glioblastoma remain poorly understood^16^. To address this gap, we first profiled intratumoral CD8^+^ T cell clonal dynamics in a recurrent glioblastoma patient who underwent initial surgical resection two weeks after a neoadjuvant ICI dose, followed by multiple subsequent ICI cycles and a second resection 17 weeks after ICI initiation (Fig. 5a). Paired scRNA-seq and TCR-seq analyses of both tumor specimens revealed a marked decrease in Shannon diversity and an increase in clonal skewing after treatment, suggesting a shift toward fewer but more dominant T cell clones following additional cycles of ICI (Fig. 5b). Notably, only around 10% of the clones detected in the tumor at timepoint 2 were present at timepoint 1, with the majority of the early clones contracting after additional cycles of treatment. This indicates substantial clonal replacement in the TME during ICI treatment (Fig. 5c). Even shared clones across the two timepoints contracted post treatment (Fig. 5c). To determine whether these new clones originated from a peripheral reservoir, we performed bulk TCRβ sequencing of matched peripheral blood samples at these time points. Many of the dominant post-treatment tumor clones were also present in the blood (circulating), supporting ICI-induced recruitment of peripheral clones into the tumor (Fig. 5d). Strikingly, predicted TRC, defined by MANAscore, were absent after a single ICI dose but emerged following multiple treatment cycles, suggesting a delayed yet specific recruitment of TRC from the periphery (Fig. 5e). At the second resection, these TRC localized primarily to the GZMK^+^ early effector and terminal effector (Term T_eff_) populations (Fig. 5e), consistent with the post-treatment TRC distribution across glioblastoma samples. Together, these findings support the idea that repeated ICI administration promotes replacement of tumor-resident clones with peripherally derived (circulating) TRC possessing effector potential.

**Fig. 5.**
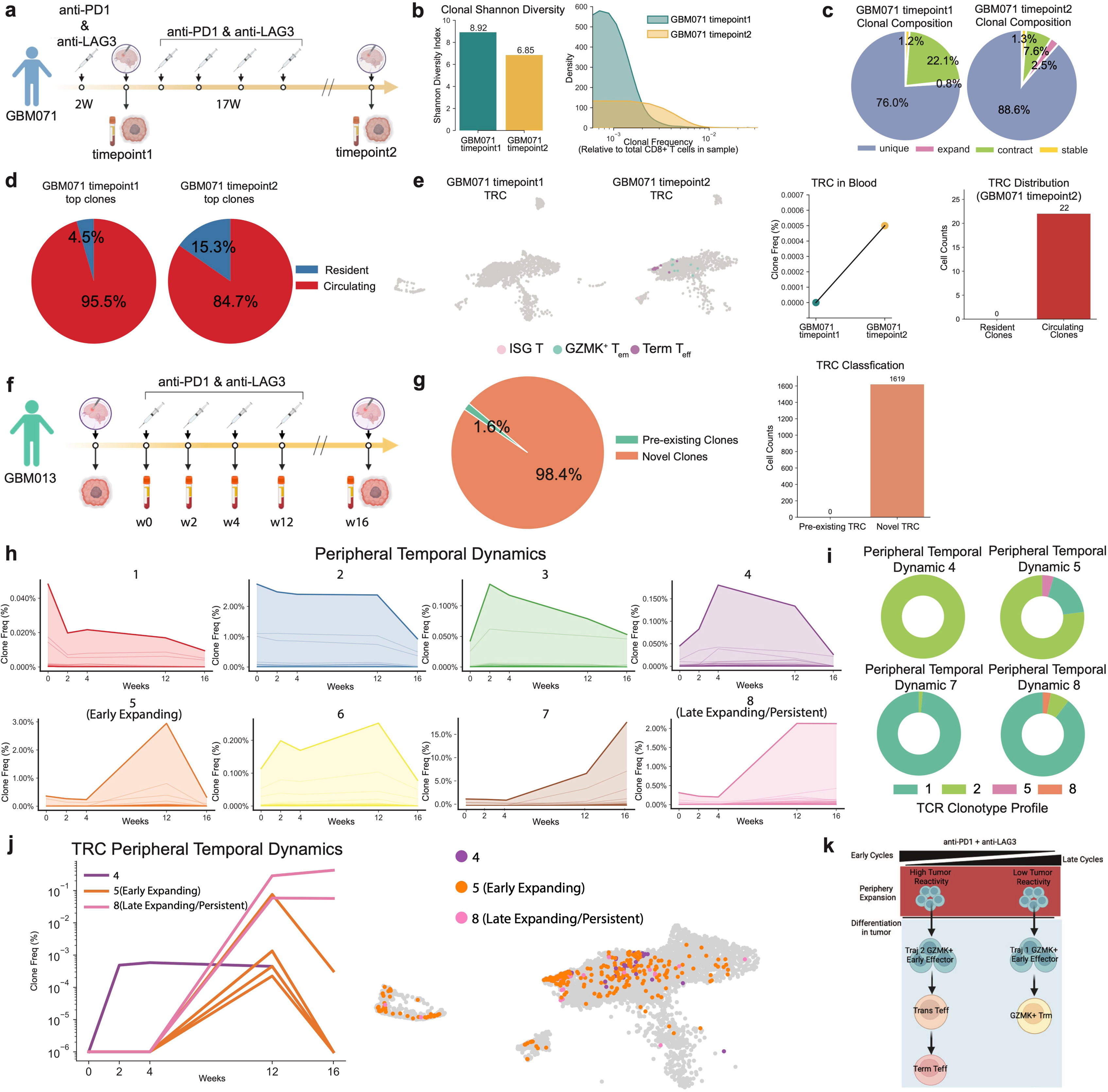
Dual ICIs lead to recruitment of novel TRC from the periphery into the tumor microenvironment. (a). Sample collection for GBM071 with paired tumor resections and peripheral blood collection, (b). Shannon diversity index (left) and clone frequency distribution (right) show reduced clonal diversity following multiple ICI cycles, (c). Composition of TILs shows dominance of unique clones after multiple ICI cycles, (d). Pie chart demonstrates expanded clones in post-treatment tumor originate primarily from the circulation, (e). Temporal tracking demonstrates that TRC emerge only after multiple ICI doses (left) and derive exclusively from the periphery (right), (f). Sample collection for GBM013, with pre- and post-treatment tumor resections and sequential peripheral blood collection, (g). Pie chart showing post-treatment tumor clones mainly consist of new clones from circulation (left). Bar chart demonstrating post-treatment tumor TRC are exclusively new clones from the periphery (right), (h). Unsupervised clustering reveals eight temporal patterns of clonal dynamics in the longitudinal peripheral blood, (i). Clonotype composition analysis demonstrates enrichment of profile/trajectory 2 cells in temporal dynamics 4 and 5 (top) and enrichment of profile/trajectory 1 cells in temporal dynamics 7 and 8 (bottom), (j). Line plot and UMAP projection demonstrate TRC mainly exhibit early expanding temporal dynamic 5 and later expanding/persistent dynamic 8. (k). Schematic illustrating GZMK^+^ early effector recruitment and differentiation. ICI induces GZMK^+^early effector cells toward a tumor-reactive effector pathway, but this response is not maintained in patients who ultimately fail to develop a therapeutic response. Instead, persistent peripheral clones diverge toward GZMK^+^ resident memory states, which are not effective at eliminating tumors directly. Ultimately, rebalancing GZMK^+^early effector differentiation to maintain a constant population of tumor-reactive effector cells may enhance ICI responsiveness.

We expanded this analysis to a second patient treated with dual ICIs, for whom we had access to tumor tissue prior to therapy, serial peripheral blood samples during treatment, and post-treatment tumor specimen obtained at the time of progression (Fig. 5f). Bulk TCRβ and scRNA/TCR-seq analyses again showed minimal overlap between pre- and post-treatment tumor infiltrating clones with only 1.6% of the clones persisted, and none of the dominant post-treatment TCR clones detectable in the baseline tumor (Fig. 5g). We next tracked the temporal dynamics of peripheral clones that appeared in the post-treatment tumor by identifying shared clones between peripheral blood and post-treatment tumor and clustering their frequency dynamic patterns across five blood draws (Extended Data Fig. 5a). This unsupervised analysis revealed eight distinct peripheral temporal dynamic patterns (Fig. 5h). These fell into three broad categories: (i) clones present at baseline (pre-treatment peripheral blood) that contract steadily during therapy (patterns 1 and 2); (ii) clones showing successive transient bursts of clonal expansion following individual ICI cycles (patterns 3, 4, and 5); and (iii) clones that began expanding around week 4 after ICI initiation and persisted or continued to expand through the last time point evaluated (4 months after treatment) (patterns 7 and 8).

To determine whether these peripheral dynamic patterns were linked to specific intratumoral differentiation fates, we analyzed the association between specific dynamic patterns with clonotrace trajectories. We found a striking dichotomy. Clones with transient bursts at specific treatment cycles (dynamic patterns 4 and 5) were almost exclusively assigned to profile/trajectory 2, differentiating from GZMK^+^ early effectors and ending as terminal effectors in the tumor. In contrast, clones that persisted through the 4-month analysis period after initial expansion (dynamic patterns 7 and 8) overwhelmingly followed profile/trajectory 1, progressing from GZMK^+^ early effectors to GZMK^+^ T_rm_ (Fig. 5i, Extended Data Fig. 5b). We then specifically examined the peripheral dynamics of TRC. We found that the majority of the TRC followed either early expanding dynamic pattern 5 (peaking at cycle 4) or late expanding/persistent dynamic pattern 8 (peaking at 12 weeks and persisting) (Fig. 5j). These findings suggest that dual ICI induces peripheral expansion of GZMK^+^ early effector TRC, which can enter the tumor and differentiate into transitional (Trans T_eff_) and terminal effector (Term T_eff_) cell states. However, this influx appears non sustained. Persistent peripheral clones instead differentiate toward resident memory fates or remain arrested in the GZMK^+^ early effector state, failing to become terminal effector TRC. This divergence in clonal fate may represent a key mechanism of resistance to ICI in glioblastoma – whereby early TRC recruitment and effector differentiation occur but are not maintained, and long-lived peripheral clones preferentially adopt non-tumor-reactive or T_rm_ phenotypes within the tumor (Fig. 5k).

## Discussion

We show here that, in contrast to the view that glioblastoma is an “immunologically cold” and immunotherapy nonresponsive cancer, combination ICI with anti-PD-1 and anti-LAG-3 induces a significant infiltration of total CD8^+^ TIL, and specifically, robust clonal expansion of tumor-reactive clones. Through the application of a new clonal embedding framework, Clonotrace, integrated with a recently described predictive algorithm for TRC, we show that dual ICI induces influx of TRC from the periphery that differentiate from a dominant GZMK^+^ population of T effector memory cells through a transitional effector state with high cytotoxic potential and ultimately to a terminal effector state with a strong exhaustion program characterized by high expression of multiple checkpoints and lower cytotoxic potential.

A central insight from our work is the delineation of a previously uncharacterized transitional effector T cell population that serves as the critical precursor to terminal effector and dysfunctional fates. This population exhibits high metabolic activity, upregulated interferon signaling, and elevated cytotoxic gene expression, including granzymes and perforin, suggesting a window of functional competence. Additionally, its unique transcriptional plasticity may render GZMK⁺ T_em_ cells particularly responsive to immunomodulatory therapies—such as checkpoint inhibitors or cytokine-based strategies—capable of redirecting their fate. Thus, the transcriptional heterogeneity and modular overlap within the GZMK⁺ Tem population highlight its central role in orchestrating the balance between activation, effector function, and exhaustion in the TME. Therapeutically, these cells appear to represent a reprogrammable reservoir of tumor-specific T cells with the potential to be reinvigorated under the right immunologic context. Importantly, these cells are clonally expanded, enriched for tumor-reactive TCRs, and transcriptionally distinct from both early effector precursors and terminal exhausted cells. This intermediate state may represent a putative therapeutic target for strategies aimed at stabilizing effector function while preventing exhaustion in glioblastoma.

Our finding that the predominant CD8 TIL population in glioblastoma expresses high levels of GZMK are consistent with the observations by Wang et al.^13^, who showed that GZMK⁺ T cells are the dominant clonally expanded CD8⁺ population in untreated glioblastoma yet display incomplete cytotoxic differentiation and a relative lack of canonical exhaustion gene expression. While their work highlights the restricted T cell repertoire and a limited activation state of TILs in untreated glioblastoma, our study expands on this foundational understanding and reveals that these GZMK⁺ T_em_ cells are the dominant population enriched for TRC in untreated glioblastoma exhibiting developmental plasticity, giving rise to both GZMK^+^ resident memory and terminal effector states. Our study further demonstrates that dual ICI can reshape this T cell landscape and repertoire. Here, we present the first comprehensive single-cell multiomic analysis that delineates the transcriptional and clonal reprogramming of CD8⁺ T cells in glioblastoma following dual blockade of PD-1 and LAG-3. Notably, dual ICI treatment drives these early effector cells into distinct differentiation trajectories, with a marked shift toward a terminal effector state marked by high checkpoint expression and regulon activation—features associated with T cell exhaustion. This suggests that dual blockade promotes an initial wave of activation, but may also accelerate terminal differentiation, potentially limiting long-term functional persistence.

We further uncover a previously unrecognized immunological phenomenon: ICI with anti–PD-1 and anti–LAG-3 not only expands these GZMK+ T cells, but also reprograms the glioblastoma-infiltrating CD8⁺ T cell compartment by inducing clonal replacement with peripherally derived TRC. These TRC enter the TME following ICI, undergo transient clonal expansion, and initiate a differentiation process toward terminal effector phenotypes along a gradient of dysfunction. Our study is among the first to demonstrate that clonal replacement in glioblastoma, driven by repeated cycles of ICI, results in the recruitment of new TRC from the peripheral blood. While similar patterns of clonal replacement have been reported in other cancers^43,44^, none of these studies distinguished TRC from the broader TIL population, which is largely composed of non-tumor-reactive bystanders. Our analysis shows that in glioblastoma, these recruited clones are tumor-reactive and follow distinct, trackable differentiation paths. These findings suggest that, in glioblastoma, peripheral T cell recruitment plays a more critical role than expansion of pre-existing tumor clones in driving responses to immunotherapy.

However, despite transiently expanding and infiltrating the tumor, the recruited TRC fail to persist in the patients with accessible tumor post-treatment, with many arrested in early differentiation or diverted toward an exhausted dysfunctional program. These findings may help explain the limited durability of response to checkpoint inhibitors in glioblastoma and underscore the challenge of sustaining effector function within an immunosuppressive microenvironment. Our findings therefore provide critical context to the impact of dual ICIs on glioblastoma T cell repertoire; namely that TRC are restrained in a GZMK⁺ progenitor state characterized by low exhaustion/checkpoint programs but also limited cytotoxic potential, potentially driven by the immunosuppressive glioblastoma microenvironment, yet can be induced to differentiate upon combination ICI treatment. A limitation of our analysis is that patients on the anti-PD-1/LAG-3 trial whose tumors responded clinically were not available for re-sampling per protocol and thus the only post-treatment initiation tumor material available for analysis were patients who relapsed and had repeat resections. It is possible that responding patients harbored a more expanded and stable pool of TRC in the GZMK^+^ early effector state, serving as reservoirs that, upon ICI treatment, actively “feed” into a transitional effector program (Trans T_eff_) marked by high cytotoxicity and relatively low checkpoint expression. This raises important therapeutic considerations on strategies for sustaining T cell persistence and function, as well as for understanding resistance mechanisms within the TME that drive divergent fates.

The bifurcation of TRC differentiation trajectories—toward terminal effector versus resident memory fates—appears to be determined early, as reflected in differential transcriptional programs within GZMK⁺ early effector cells. Inhibitory receptors and regulons were differentially expressed among early GZMK^+^ precursors with divergent trajectories, suggesting that early lineage decisions post-ICIs are tightly linked to the eventual fate and function of TILs. Importantly, we find that the transient effector window—encompassing the Trans T_eff_ populations—is characterized by high cytotoxic gene expression, metabolic activity, and chemotactic signaling. This suggests a metabolically and functionally poised state that may be particularly amenable to ICI reinvigoration. We propose that this immune response and cellular state are not maintained in patients who ultimately fail to develop a therapeutic response. Therapeutic strategies that seek to stabilize this transitional state—such as cytokine supplementation, metabolic reprogramming, or epigenetic modulation—could potentially extend the therapeutic window of ICI and promote durable anti-tumor immunity.

Taken together, our work significantly advances the understanding of the glioblastoma T cell repertoire by showing that dual ICI reshapes clonal dynamics and T cell fate in ways not previously appreciated by promoting the influx and reprogramming of tumor-reactive CD8⁺ T cells along a unique, clonally traceable differentiation axis in glioblastoma. It underscores the importance of clonal recruitment from the periphery and highlights the need to preserve early effector T cell potential within the TME. These insights argue for the rational design of therapeutic combinations that go beyond checkpoint inhibition alone—potentially incorporating therapies that preserve early effector function and target metabolic vulnerabilities within the TME-to sustain anti-tumor immunity in glioblastoma.

## Methods

### Human subjects

Patients were prospectively enrolled into this study at Johns Hopkins Hospital. Written informed consent was provided by all participants according to approved Institutional Review Board Protocol. Fresh tumor tissue and blood were collected at the time of surgery following confirmation of primary glial neoplasm on frozen section. None of the patients was treated with chemotherapy or radiation prior to tumor resection. Tumor samples span all glioma grades and subtypes. Tumors were categorized according to the 2021 WHO Classification of Central Nervous System Tumors based on the combination of relevant histopathologic and molecular features. Patients undergoing anterior temporal lobectomy for seizure control were consented and non-neoplastic tissue from anterior temporal lobectomy resection was collected as control.

### Tumor dissociation

Patient tumors or non-malignant brain tissue were mechanically cut into about 1mm^3^ tissue in DMEM medium (Gibco). They were then enzymatically digested with MACS tumor dissociation kit (Miltenyi Biotec) on the gentleMACS Dissociator (Miltenyi Biotec) per manufacture instructions. Dissociated cells were filtered by a 70-mm strainer and centrifuged at 300g for 7 min. After removing the supernatant, the pelleted cells were suspended in red blood cell lysis buffer (Miltenyi Biotec) for 2 min to remove red blood cells. Tumor samples with significant extent of myelin underwent myelin removal process using magnetic Myelin Removal Beads (Miltenyi Biotec) on the autoMACS (Miltenyi Biotec). The cells were then washed with sorting buffer (PBS supplemented with 2% FBS) and accessed for viability using trypan blue exclusion.

### Peripheral blood mononuclear cell collection

Peripheral blood mononuclear cells (PBMCs) were isolated after Ficoll-Paque centrifugation. Briefly fresh peripheral blood was collected at the time of surgery in EDTA anticoagulant tubes and layered on top of Ficoll-Paque after twofold dilution with phosphate-buffered saline (PBS). After centrifugation (2200 rpm, 20 minutes, no brake), PBMCs were collecting from the mononuclear cell band layer. PBMCs were viability frozen in 10 million per mL until future use.

### Fluorescence-activated cell sorting

Single cell suspensions of the tumor and non-malignant brain tissue were stained with antibodies for 30 minutes at 4°C against CD45+ (PE, BD Bioscience), CD3+ (APC, BD Bioscience) and nuclear stain DyeCycle Violet (Invitrogen) for FACS sorting, performed on Beckman Coulter MoFlo XDP. Cells were sorted into three live populations, T cells (CD45+CD3+), non-T immune cells (CD45+CD3-), and non-immune cells (CD45-, DyeCycle Violet+) directly into cold PBS + 1% bovine serum albumin (BSA, Sigma-Aldrich) with a post-sort purity of >98%. Sorted cells were then counted and assessed for viability using trypan blue and re-suspended in 700cells/ml in PBS + 0.04% BSA.

### Single-cell RNA and TCR sequencing and processing

scRNA-seq and scTCR-seq of T cells were performed using the 10X Single Cell 5’ Immune Profiling Kit v1.1 (10X Genomics, Pleasanton, CA, USA). Cells were captured in droplets at a targeted cell recovery of 5000-10,000 cells per lane. scRNA library generation was performed per protocol. Following cell barcoding in droplets and reverse transcription, emulsions were broken and cDNA purified using Dynabeads. cDNA was amplified by 13 PCR cycles for 5’ immune profiling. Amplified cDNA was then used for 5’ gene expression library construction and TCR enrichment. Sequencing was performed using an Illumina NovaSeq 6000 with 310 million reads per sample and a sequencing configuration of 26x8x98 (UMI × Index × Transcript read). The Cell Ranger 3.0.2. pipeline software (10X genomics, California) was used to align reads and generate expression matrices for downstream analysis.

#### ScRNA-seq filtering and normalization

After combining raw counts from the filtered output of CellRanger, we further filtered cells based on both minimum number of genes and counts. Selection of each was performed using histograms of each parameter by cell. In each case, a minimum threshold of 500 genes and 750 UMI counts was sufficient to exclude populations of low-quality cells visible as the lower portion of the bimodal/multimodal distributions visualized in the histogram. Genes expressed in fewer than .1% of the cells were also removed. Finally, counts were normalized to the total UMI count by cell and log-scaled using Seurat.

#### Data integration

Data integration was performed using Seurat v3 using reciprocal principal component analysis to project cells into a shared space followed by identification of anchors for mutual nearest neighbor integration. One female and one male patient were selected as references for integration.

#### Scaling, principal component analysis, clustering, and dimensional reduction

Scaling of both corrected and uncorrected normalized gene expression values was calculated, although the uncorrected values were used only for visualization of differential gene expression with heatmaps. In both cases, values were scaled to a mean of zero and a standard deviation of one by gene. The scaled, corrected values were used for principal component analysis, after which 60 principal components were selected based on elbow plot for all data sets. The 60 principal components were used for dimensional reduction by uniform manifold approximation projection (UMAP) as well as generation of a shared-nearest neighbor network followed by Louvain clustering. Clusters with very similar gene expression profiles were combined and, in some cases, clusters were isolated for further clustering.

#### Differential gene expression

To identify marker genes distinguishing cell populations, we performed differential expression analysis using the Wilcoxon rank-sum test, as implemented in Scanpy’s rank_genes_groups function. Genes were considered significant if they met all the following criteria: adjusted *p-value* < 0.05, log fold-change > 0.25, and expression in more than 30% of cells within the group of interest.

#### Cell cycle scoring

Seurat’s CellCycleScoring was used to identify cells with high G2M or S phase contributions. While this was not used for regression of gene expression values during scaling of all cells, it was used to identify clusters of cells undergoing mitosis. These clusters were isolated and then scaling was performed on these cells alone with regression of cell cycle scores. Finally, principal component analysis and clustering was performed on the regressed, scaled values to attempt to identify types of cells present in the cycling clusters.

##### Hotspot module analysis and Gene Set enrichment analysis

Hotspot module analysis was performed on raw count matrices using its default negative-binomial variance model^24^. A K nearest neighborhood graph was constructed with k = 30. The first 15 principal components were used as Hotspot’s transcriptomic latent space, and the top 3,000 genes (false discovery rate < 0.05 across the dataset) were retained. These genes were grouped into co-regulatory modules via agglomerative clustering, enforcing a minimum module size of 150 genes. For each resulting module, over-representation analysis was conducted using GSEApy’s^45^ Enrichr interface against the KEGG_2019_Human, GO_Biological_Process_2018, and Reactome_2016 libraries; the Benjamini–Hochberg–adjusted p value was used to assess pathway enrichment.

##### Viral Specific and Tumor Specific Geneset Collection and Scoring

Gene sets for CD8^+^ tumor reactivity from multiple previously published models were collected and merged into a single tumor reactivity signature^32,46,47^. A viral-specific signature was obtained from a published viral reactivity model^25^. Module scores for each signature were calculated on the scaled, normalized expression matrix using Scanpy’s score_genes function^48^.

##### MANAscore and TRC identification

MANAscore was computed for each CD8^+^ T cell using both imputed and non-imputed data via the MANAscore package^33^. For each sample, a high-score threshold was defined at the stabilization point of the long-tail distribution in both imputed and non-imputed MANAscore values. Cells exceeding both thresholds were classified as MANAscore-high. Clonotypes containing more than two MANAscore-high cells were designated predicted tumor-reactive cells (TRC).

##### Expansion index

The expansion index quantifies the extent of clonal expansion in a T cell population. It is designed to depend solely on the number of cells per clone. If a clone consists of only one cell, it is considered unexpanded, and its contribution to the expansion index is zero. In contrast, clones with multiple cells are treated as expanded, and their contribution increases with clone size.

Based on this intuition, we define the expansion index using the cell frequency of each clone. For a sample with total cell count *p*, assume there are *m* clonotypes (clones) within a given cell type *t*. Let *n_it_* be the number of cells assigned to clone *i* in cell type *t*. Then the expansion index for cell type *t* is defined as:

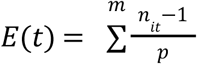

This formulation ensures that:

- Clones with only one cell do not contribute to the index.
- The index increases with both the number and size of expanded clones.
- The value is normalized by the total number of cells, allowing for comparison across samples.

##### Pseudobulk gene expression analysis

DecoupleR package is being applied for pseudobulk analysis^49^. GZMK^+^ Tem cells were aggregated into pseudobulk profiles by summing raw counts for each gene across all cells sharing a given sample. The resulting gene by pseudobulk matrix was then normalized for library size using the TMM method implemented in edgeR, and low-expression genes (fewer than 10 counts in at least two pseudobulk samples) were excluded from further analysis. Principal component analysis (PCA) was applied to the normalized expression matrix, and one-way ANOVA was used to assess whether sample scores on each principal component differed significantly across tumor grades.

##### Clonotrace

The Clonotrace package is used to perform clonotype profile analysis. Clones are defined as cell populations sharing identical CDR3α and CDR3β protein sequences. Clones with low expansion (fewer than 10 cells) are excluded from further analysis. Pairwise distances between clones are calculated in the top 30 principal components of the transcriptomic space using optimal transport and clustered with the Leiden algorithm.

To construct the clone-pruned cell UMAP, we first generate a cell k-nearest neighbor (kNN) graph from the transcriptomic PCA space. In parallel, clone-level distances are projected into a multidimensional scaling (MDS) space to build a clone kNN graph. Edges between cell neighbors whose corresponding clones are not neighbors in the clone graph are then pruned. The resulting pruned cell graph is used to compute a UMAP embedding that reflects both transcriptional similarity and clonal structure.

Profile enrichment within cell clusters is assessed via Fisher Exact Test. To identify differentially expressed genes (DEGs) between profiles within a given cluster, we identify, for each cell in the target profile, its top k (default: 3) nearest neighbors from other profiles based on pseudotime distance. A Wilcoxon rank-sum test is applied to all genes expressed in at least 30 cells across the dataset.

##### Pseudotime Analysis

The top 30 Harmony-integrated components were used as input to Palantir to construct diffusion maps. The starting cell was defined as the cell with the lowest expression of a checkpoint gene signature. Palantir was run with 1,500 waypoints and 30 nearest neighbors. Gene expression was imputed and smoothed along pseudotime using MAGIC for visualization. To identify dynamic genes, a generalized additive model (GAM) with a cubic spline was fitted to log-normalized expression values as a function of pseudotime using pygam. P values for the spline term were adjusted using the Benjamini–Hochberg method; genes with an adjusted P < 0.05 were considered significantly dynamic.

##### Bulk TCR sequencing

DNA was isolated from (i) formalin-fixed, paraffin-embedded (FFPE) tumor blocks obtained prior to combination ICIs and (ii) 5 million peripheral-blood mononuclear cells (PBMCs). Tumor cores were macro-dissected directly from the FFPE tissue, and PBMC were processed with the same extraction kit. Library preparation, ultra-deep amplification of TCR-β CDR3 regions, high-throughput sequencing, and primary repertoire analysis were performed by Adaptive Biotechnologies using the immunoSEQ platform.

##### Cyclone and TCR trajectory analysis

Cyclone was used to trace the temporal dynamics of tumor-infiltrating CD8^+^ T cells in peripheral blood^50^. Clonotypes in bulk TCR-seq were identified by matching β-chain sequences against single-cell TCR-seq data. Clonotypes exhibiting a ≥2-fold change in frequency and an absolute frequency change ≥2.5% in any two timepoints were retained for trajectory analysis. For each retained clonotype, frequencies across all timepoints were normalized via min–max scaling. Pairwise Euclidean distances between these normalized frequency vectors were computed, and agglomerative hierarchical clustering with Ward’s linkage was applied. Final clusters were defined using a maximum-cluster criterion.

##### Statistical Analysis

Statistical analyses were performed in Python using the statsannotations package. All comparisons were unpaired and used two-sided Mann Whitney U tests, with p-values adjusted for multiple testing via the Benjamini–Hochberg method. Significance levels were defined as: p < 0.05 (*), p < 0.01 (**), and p < 0.001 (***). Error bars denote the 95th percentile confidence interval.

## Data availability

Raw and processed single-cell RNA-seq and TCR-seq data will be made available at the NCBI’s Gene Expression Omnibus (GEO). All other data are available in the main text or the supplementary materials.

## Code availability

Custom computational methods beyond routine algorithms and packages are wrapped into the R package Clonotrace, publicly accessible through Github (https://github.com/yuntianf/Clonotrace)

## Acknowledgements

We thank Jennfer Meyers, Kornel Schuebel, Anuj Gupta, Alyza Skaist, and other members of the Experimental and Computational Genomics Core, supported by NCI Cancer Center Support Grant P30 CA006973

## Author contributions

Conceptualization: CJ, ML, DP, NRZ. Methodology: CJ, DP, YF, NRZ, MW, ZZ, KNS. Investigation: CJ, MW, SB. Visualization: CJ, MW, YF, YN. Funding acquisition: CJ, DP, ML, NRZ, SY. Project administration: CJ, NRZ, DP, ML. Supervision: CJ, NRZ, ML, DP. Writing – original draft: CJ, MW, DP, YF, NRZ. Writing – review & editing: MW, YF, SB, YN, DM, MZ, CL, JC, ZZ, TZ, HJ, VY, JW, CB, KNS, ML, DP, NRZ, CJ.

---

**Extended Data Fig. 1.**
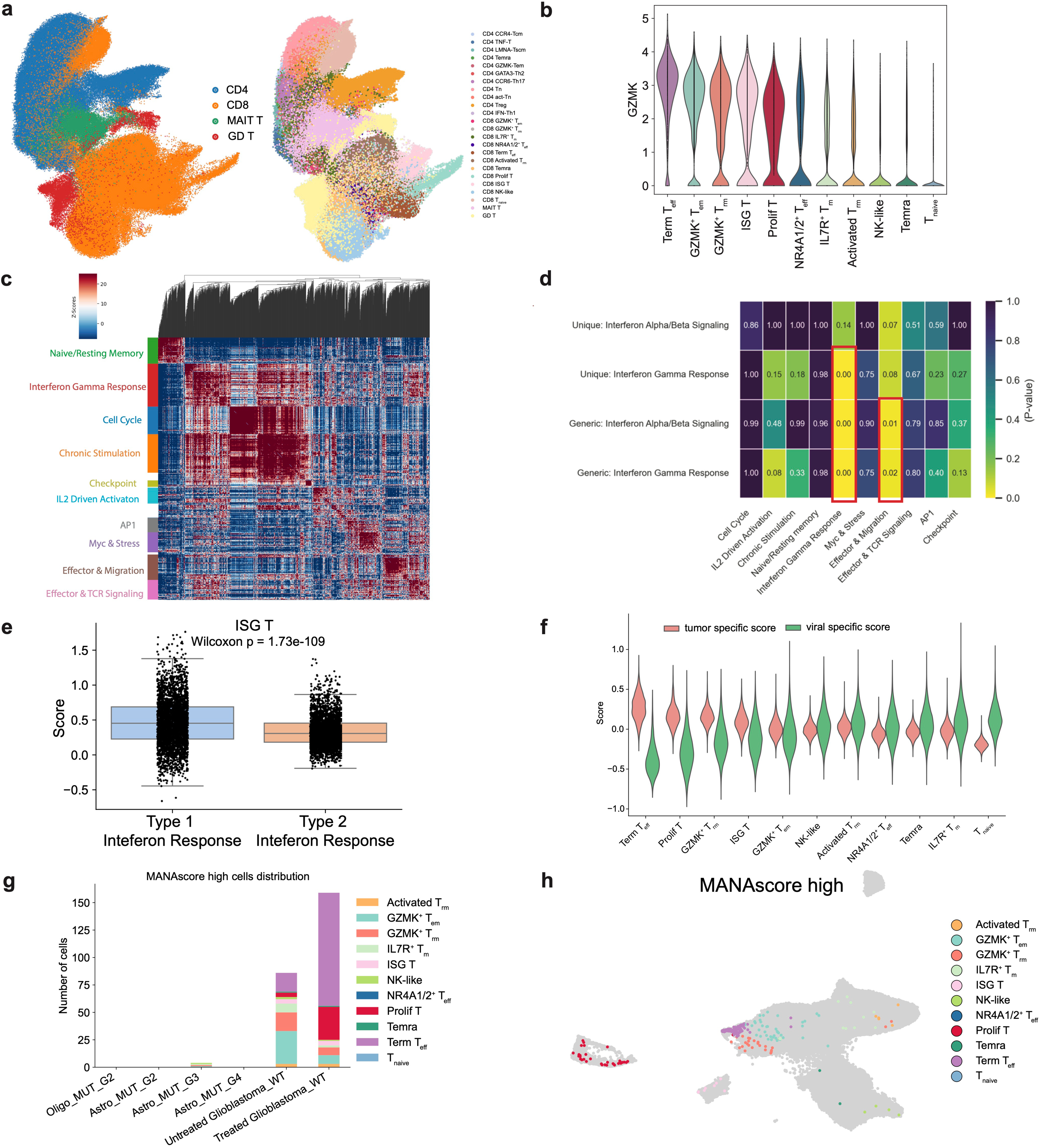
Tumor-reactive CD8^+^ T cells in glioblastoma exhibit GZMK^+^ phenotypes. (a). UMAP projection of the expression profile of all CD45^+^CD3^+^ cells reveals 11 clusters of CD8^+^ T cells, 11 clusters of CD4^+^ T cells, a γŏ T cell cluster, and a MAIT T cell cluster, (b). Violin plot demonstrating increased GZMK expression in T_em_, T_rm_, Term T_eff_, ISG T, and ProlifT clusters, (c). Heatmap demonstrating auto-correlation of top 3000 informative genes identified by Hotspot, revealing ten distinct modules, (d). Heatmap showing over-enrichment test p-values of all genes and unique genes in type I and type II interferon response across the ten modules suggesting interferon gamma response rather than interferon alpha/beta response enrichment in the interferon response module, (e). Boxplot showing that the ISG T subset exhibits a moderate bias toward type I interferon response, (f). Violin plot showing enrichment of published tumor-specific T cell gene signature score in Term T_eff_, ProlifT, GZMK^+^ T_rm_, ISG T, and GZMK^+^ T_em_. (g). Distribution of MANAscore high cells across different tumor grades demonstrates virtually absent MANAscore high cells in IDH-mutant gliomas, (h). MANAscore high cells in glioblastoma are enriched in Term T_eff_ cells.

**Extended Data Fig. 2.**
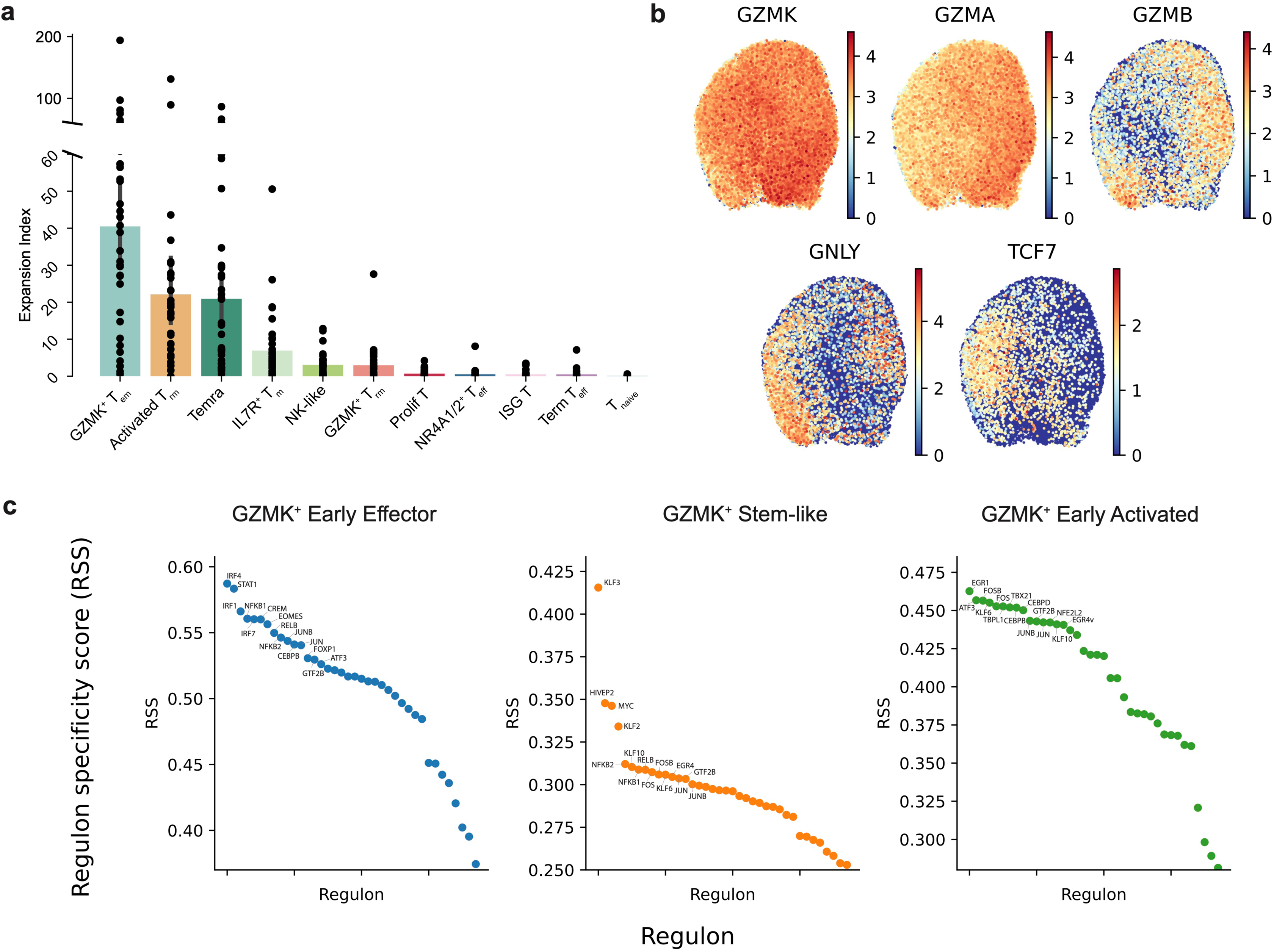
Clonally expanded GZMK÷ T_em_ demonstrate a spectrum of cell states with distinct regulon activities. (a). Expansion index of CD8÷ T cell clusters demonstrates that GZMK÷ T_em_ is the most clonally expanded population, (b). Feature plots demonstrating selective expression of cytotoxic and sternness genes across GZMK÷ T_em_ subsets, (c). Top 15 most specific regulons for the three GZMK÷ T subclusters derived from SCENIC.

**Extended Data Fig. 3.**
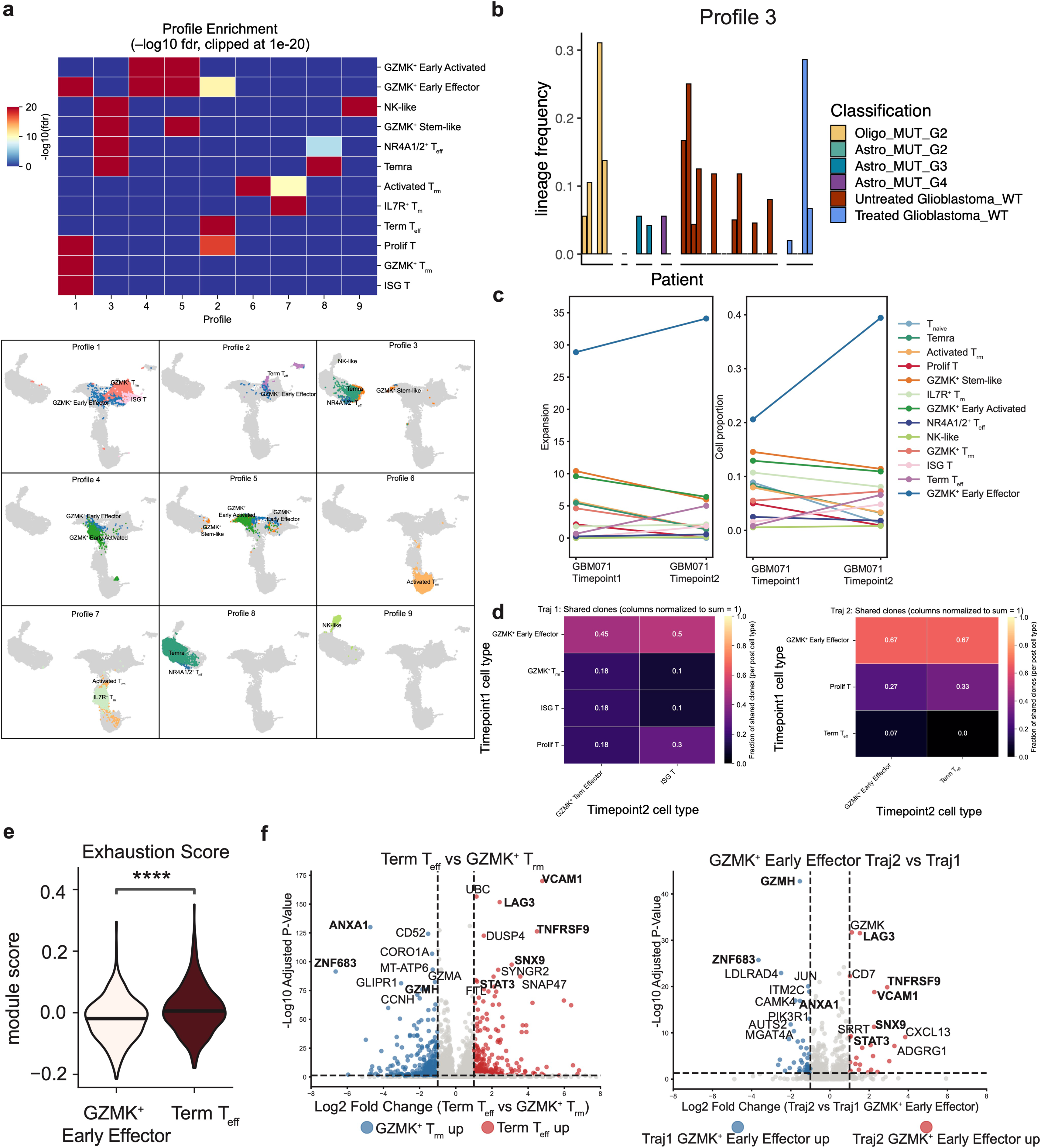
Clonotrace identifies distinct TOR clonotype profiles and reveals early differentiation divergence of GZMK^+^ early effector cells. (a). Heatmap of FDR of T cell cluster enrichment across clonotype profiles (top), only FDR < 0.05 is highlighted; Projection of 9 clonotype profiles on clonal embedding weighted UMAP in PCA space (bottom) demonstrating distribution of clonotype profiles across T cell subclusters, (b). Bar plot demonstrating the enrichment of clonotype profile 3 across all glioma groups, (c). Clonal expansion and infiltration changes of each CD8÷ T cell subset at two time points in a single patient, (d). Heatmap showing the fraction of clones from the timepointl treated state contributing to the timepoint2 treated state of the two trajectories (Traj 1: profile 1, Traj 2: profile 2). (e). Violin plot showing exhaustion module scores in GZMK÷ early effector and Term T_eff_ in Profile 2/Traj 2. (f). Volcano plot of differentially expressed genes in GZMK÷ early effector Traj2 vs Trajl and Term T_eff_ vs GZMK÷ T_rm_ with shared genes between the precursor cells and their terminal states highlighted in bold.

**Extended Data Fig. 4.**
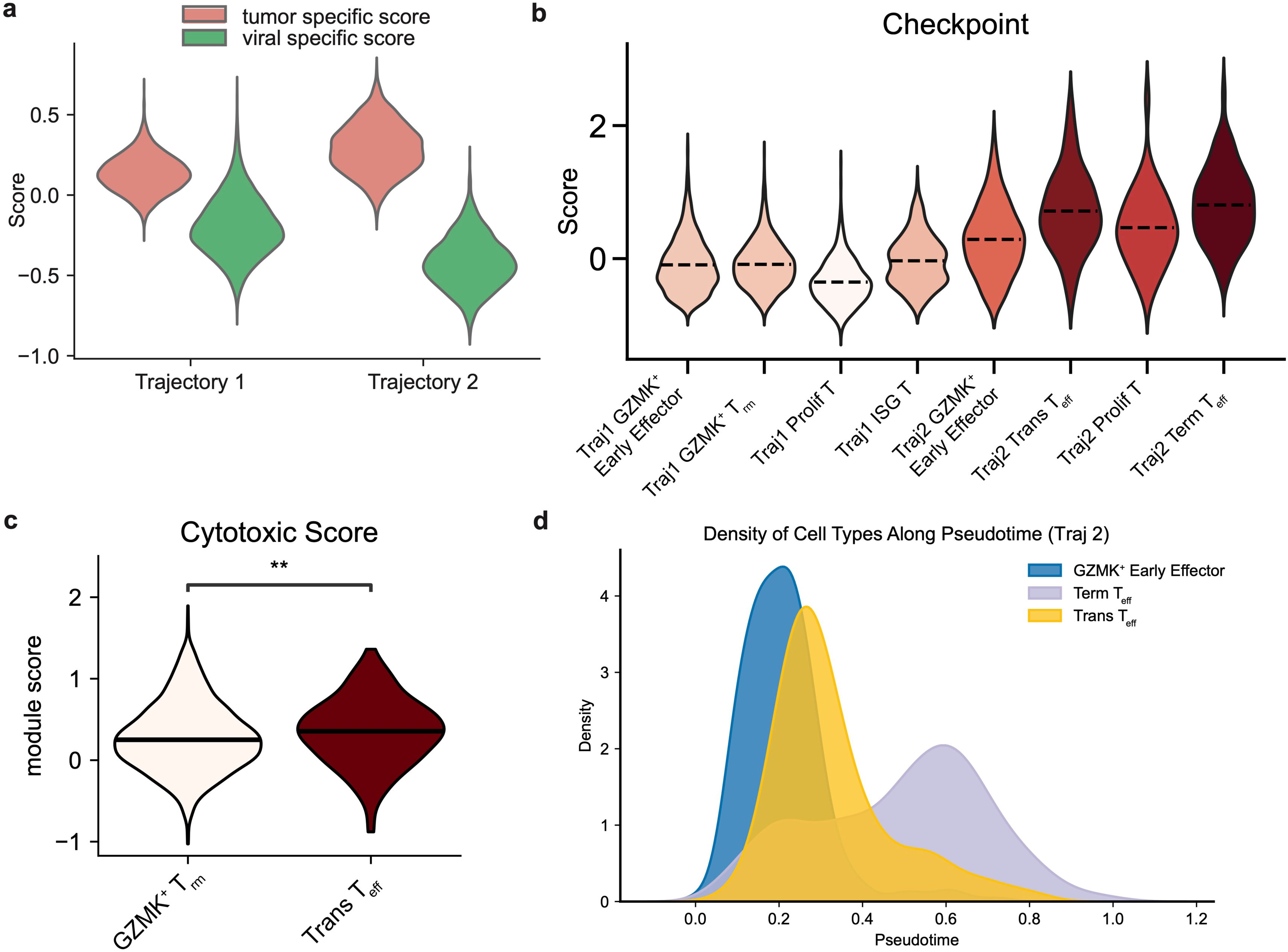
Dual checkpoint blockade drives a GZMK^+^ early effector trajectory enriched for tumor specificity, checkpoint expression, and cytotoxicity. (a). Violin plot showing increased tumor-specific and reduced viral-specific gene signature scores in Traj 2 compared to Traj 1 using published signatures, (b). Violin plot of checkpoint score across Traj 1 and Traj 2 subsets demonstrating increased checkpoint expression across Traj 2. (c). Violin plot demonstrating increased cytotoxic score in Trans T_eff_ cells compared to GZMK^+^ T_rm_ cells, (d). Density plot of Traj 2 cell subsets along diffusion map pseudotime demonstrating the temporal relationship between the three T-cell subsets.

**Extended Data Fig. 5.**
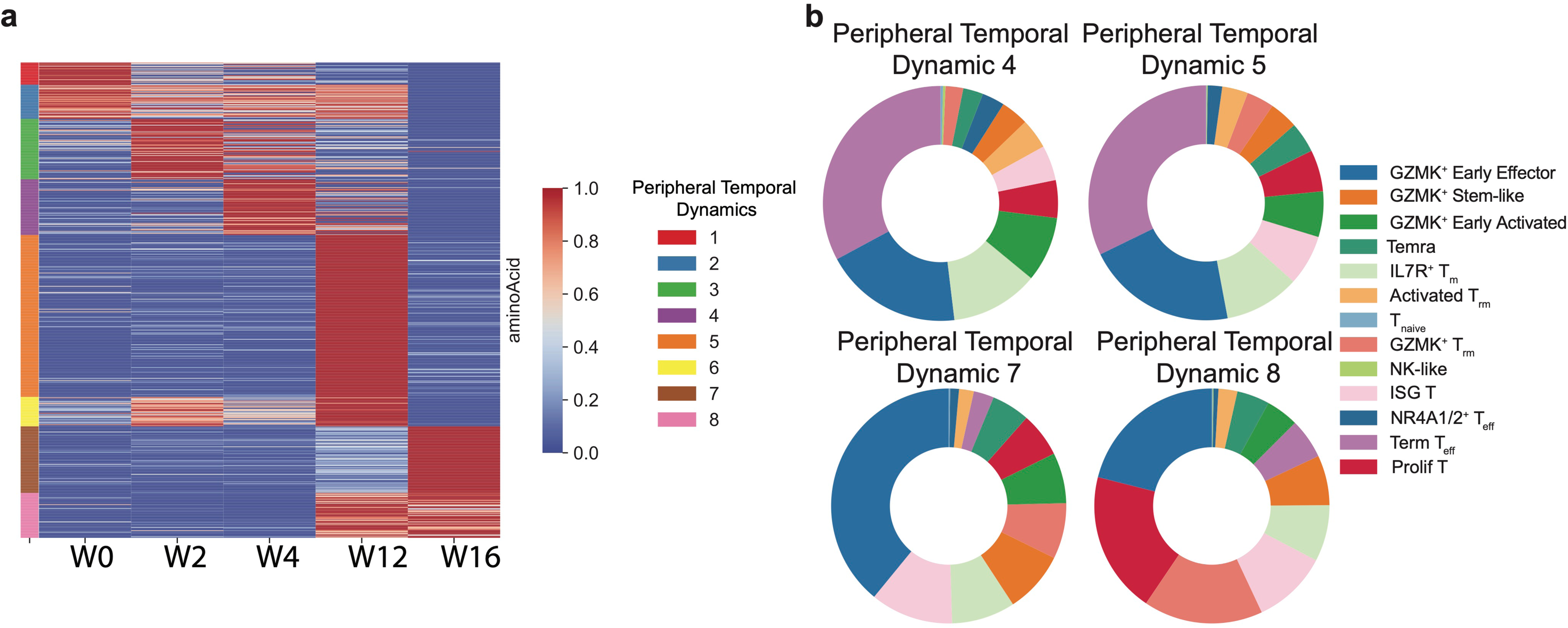
Dual ICI therapy drives temporal waves of peripheral CD8^+^ T-cell expansion. (a). Heatmap depicting peripheral blood clonal frequency changes of CD8^+^ T cell clones detected in the tumor at the time of progression (week 16) over the course of treatment, (b). Cell cluster composition of peripheral temporal dynamics shows increased Term T_eff_ in the early expanded cycles (temporal dynamics 4 and 5) and GZMK^+^ early effector subset in the late expanded cycles (temporal dynamics 7 and 8).

