## Supplemental Table 1 and 2 for "Dual checkpoint blockade of glioblastoma with Anti-PD-1 and Anti-LAG-3 promotes expansion of tumor-reactive T cell clones along a unique pathway of differentiation"

**Extended Data Table 1. Patient clinical characteristic and sequencing statistics**

| Patient ID | Age/Sex | 2021 Classification | WHO Grade | CD45+CD3+ |
|---|---|---|---|---|
| GBM006 | 67/F | Glioblastoma, IDH-WT | 4 | 3652 |
| GBM009 | 70/M | Glioblastoma, IDH-WT | 4 | 2912 |
| GBM010 | 53/M | Glioblastoma, IDH-WT | 4 | 5829 |
| GBM013 | 52/M | Glioblastoma, IDH-WT | 4 | 11105 |
| GBM029 | 74/F | Glioblastoma, IDH-WT | 4 | 2805 |
| GBM030 | 66/M | Glioblastoma, IDH-WT | 4 | 5546 |
| GBM035 | 66/F | Glioblastoma, IDH-WT | 4 | 2415 |
| GBM036 | 61/F | Glioblastoma, IDH-WT | 4 | 6800 |
| GBM037 | 73/F | Glioblastoma, IDH-WT | 4 | 6375 |
| GBM040 | 61/F | Glioblastoma, IDH-WT | 4 | 1838 |
| GBM043 | 63/M | Glioblastoma, IDH-WT | 4 | 7102 |
| GBM045 | 44/F | Oligo, IDH-mutant & 1p/19q codel. | 2 | 13018 |
| GBM046 | 64/M | Glioblastoma, IDH-WT | 4 | 1735 |
| GBM048 | 56/F | Glioblastoma, IDH-WT | 4 | 11500 |
| GBM049 | 71/M | Glioblastoma, IDH-WT | 4 | 11423 |
| GBM050 | 64/F | Glioblastoma, IDH-WT | 4 | 12818 |
| GBM051 | 66/M | Glioblastoma, IDH-WT | 4 | 21456 |
| GBM052 | 40/M | Glioblastoma, IDH-WT | 4 | 15090 |
| GBM054 | 64/F | Glioblastoma, IDH-WT | 4 | 9542 |
| GBM055 | 37/F | Astrocytoma, IDH-mutant | 3 | 15431 |
| GBM056 | 78/F | Glioblastoma, IDH-WT | 4 | 12700 |
| GBM057 | 47/F | Glioblastoma, IDH-WT | 4 | 17583 |
| GBM059 | 38/F | Oligo, IDH-mutant & 1p/19q codel. | 2 | 7432 |
| GBM060 | 31/F | Astrocytoma, IDH-mutant | 3 | 4776 |
| GBM062 | 51/F | Glioblastoma, IDH-WT | 4 | 6524 |
| GBM064 | 28/M | Astrocytoma, IDH-mutant | 4 | 12668 |
| GBM065 | 77/M | Glioblastoma, IDH-WT | 4 | 14104 |
| GBM066 | 55/M | Glioblastoma, IDH-WT | 4 | 17726 |
| GBM068 | 42/F | Oligo, IDH-mutant & 1p/19q codel. | 2 | 4280 |
| GBM069 | 66/F | Oligo, IDH-mutant & 1p/19q codel. | 2 | 11583 |
| GBM070 | 32/F | Astrocytoma, IDH-mutant | 2 | 2790 |
| GBM071 | 42/M | Glioblastoma, IDH-WT | 4 | 5938 |
| GBM073 | 33/M | Astrocytoma, IDH-mutant | 4 | 15246 |
| GBM074 | 64/F | Oligo, IDH-mutant & 1p/19q codel. | 2 | 17912 |
| GBM075 | 31/F | Astrocytoma, IDH-mutant | 3 | 11250 |
| GBM081 | 24/F | Oligo, IDH-mutant & 1p/19q codel. | 2 | 6819 |
| GBM082 | 83/F | Glioblastoma, IDH-WT | 4 | 5596 |
| GBM087 | 42/M | Glioblastoma, IDH-WT | 4 | 4089 |

Oligo: oligodendroglioma, codel: co-deleted, WT: wildtype

**Extended Data Table 2. Tumor-specific and viral-specific transcriptional signatures**

| **Signature** | **NeoTCR-CD8** | **Tumor Specific** | **MANA-TIL** |
|---|---|---|---|
| **Reference** | **Lowery, Rosenberg, Science 2022** | **Oliviera, Wu, *Nature* 2021** | **Caushi, Smith, *Nature* 2021** |
| | ATP10D | KRT86 | HLA-DPB1 |
| | GZMB | RDH10 | IFNG |
| | ENTPD1 | TYMS | MIR4435-2HG |
| | KIR2DL4 | HMOX1 | TNS3 |
| | LAYN | GNG4 | HLA-DPA1 |
| | HTRA1 | CXCL13 | ENTPD1 |
| | CD70 | AFAP1L2 | GEM |
| | CXCR6 | ACP5 | GZMA |
| | HMOX1 | MYO1E | CCL3 |
| | ADGRG1 | LAYN | HLA-DQB1 |
| | LRRN3 | TNS3 | HLA-DRB1 |
| | ACP5 | TNFSF4 | HLA-DQA1 |
| | CTSW | AKAP5 | HLA-DRB5 |
| | GALNT2 | HAVCR2 | HLA-DQA1 |
| | LINC01480 | ENTPD1 | HLA-DRB5 |
| | CARS | SLC2A8 | HLA-DRA |
| | LAG3 | AC243829.4 | CXCL13 |
| | TOX | ZBED2 | |
| | PTPRCAP | MCM5 | |
| | ASB2 | CAV1 | |
| | ITGB7 | GOLIM4 | |
| | PTMS | TRAV21 | |
| | CD8A | VCAM1 | |
| | GPR68 | PON2 | |
| | NSMCE1 | MTSS1 | |
| | ABI3 | CD38 | |
| | SLC1A4 | TRBV11-2 | |
| | PLEKHF1 | MS4A6A | |
| | CD8B | TOX2 | |
| | LINC01871 | CSF1 | |
| | CCL4 | GALNT2 | |
| | NKG7 | FXYD2 | |
| | CLIC3 | PLPP1 | |
| | NDFIP2 | LMCD1 | |
| | PLPP1 | MYL6B | |
| | PCED1B | LAG3 | |
| | CXCL13 | HLA-DRA | |

PDCD1
PRF1
HLA-DMA
GPR25
CD9
TIGIT
HLA-DRB5
SYTL3
SLF1
NEK1
CASP1
SMC4
TSEN54
PLSCR1
GNPTAB
HLA-DPB1
PLEKHA1
ARHGAP9
ALOX5AP
SH3BP1
NCF4
NELL2
GATA3
PPM1M
TNFRSF1A
AC022706.1
MCM5
HLA-DRB1
TNFSF10
TRIM21
HDLBP
ERN1
CALHM2
SASH3
ACTA2
MAST4
CAPG
MPST
IGFLR1
GZMA
CD27
ITGAE
SLA2

IGFLR1
CCDC50
CD27
KIAA1324
CDKN2A
CD70
ABHD6
CTLA4
PDCD1
GEM
NUSAP1
TOX
CXCR6
NMB
HOPX
CLIC3
INPP5F
SNAP47
TSHZ2
HLA-DMA
SIT1
HLA-DRB1
TUBB
PYCARD
ADGRG1
HLA-DQA1
PRF1
HLA-DPA1
PTMS
CKS1B
HIPK2
CHST12
LSP1
FAM3C
SLC1A4
NUDT1
DNPH1

RHOC
COMMD8
MYO1G
SP140
PHPT1
CD2BP2
PLEKHO1
STAM
MRPL16
IL2RB
ID2
TESPA1
GOLGA8B
MIS18BP1
VAMP5
DAPK2
HLA-DPA1
TSG101
IL4R
CCND2
CTSC
TRAF3IP3
NLRC3
ORAI3
GNLY
MIR155HG
CARD16
CD82
ECH1
JAML
EEF1G
ETFB
DAXX
RBM4
HCST
RAB27A
YPEL2
CHST12
ARPC1B
PDIA4
PDIA6
AC243960.1
TBC1D10C

PTPN6
PYCARD
BST2
BTN3A2
MTG1
MLEC
DUSP4
GSDMD
SLAMF1
IFI6
PCID2
GIMAP1
ITGA1
CSNK2B
CDK2AP2
MYO1F
AC004687.1
PTTG1
APOBEC3C
TSPAN14
MOB3A
STXBP2
LCP2
PLA2G16
LINC00649
CST7
TADA3
SIT1
APOBEC3G
SUSD3
CD3G
CCL5
CDC25B
TNFRSF1B
HMGN3
THEMIS
ASF1A
CTNNB1
FIBP
CCDC85B
POLR3GL
GIMAP6
ARL6IP1

CALCOCO2
CCPG1
KLRB1
ACAA2
ISG15
EIF4A1
CAT
MANF
XAB2
GRINA
GLO1
LSM2
SLFN5
FKBP1A
AKNA
TAP1
LMO4
APEH
C12orf75
TMEM14A
DNPH1
C17orf49
NUDT5
MGAT1
CCDC69
EIF4EBP1
PDHB
ARL3
UCP2
IFI35
HSBP1
LYST
MRFAP1L1
ITGAL
AIP
RASAL3
CAPN1
ITGB1
RBPJ
LBH
DYNLL1
NME2
MT1F

SYNGR2
ABTB1
ZGPAT
CD63
ILK
SKA2
TMEM204
ACO2
HOPX
CRIP1
OXNAD1
CCS
GRAP2
GSTO1
HADHB
IL16
PIN4
CUEDC2
CALM3
SAMSN1
HM13
SNAP23
LPCAT4
FAAP20
EFHD2
PRDX3
CCM2
C22orf39
SDHA
ARRDC1
MAP4K1
NDUFA13
IL27RA
C14orf119

| **Virus Specific** | **Influenza TIL** |
|---|---|
| **Oliviera, Wu, *Nature* 2021** | **Caushi, Smith, *Nature* 2021** |
| RARA | IL7R |
| GADD45B | GPR183 |
| C1orf21 | KLRC1 |
| AOAH | GPR15 |
| MATK | ARL4A |
| SATB1 | PTGER2 |
| MBP | AUTS2 |
| ANTXR2 | RARA |
| RORA | PIK3R1 |
| CCR7 | LYAR |
| ANXA1 | NUDT11 |
| BACH2 | KLF3 |
| GLUL | CCND3 |
| TNFSF14 | CLDND1 |
| AUTS2 | YPEL5 |
| PERP | |
| EPHA4 | |
| TCF7 | |
| SELL | |
| MYC | |
| IL7R | |
| CD300A | |
| ITGA5 | |
| GPR183 | |
| KLF3 | |
| S1PR1 | |
